# How do antimicrobial peptides interact with the outer membrane of Gram-negative bacteria? Role of lipopolysaccharides in the peptide binding, anchoring and penetration

**DOI:** 10.1101/2023.08.30.555525

**Authors:** Justus C. Stephani, Luca Gerhards, Bishoy Khairalla, Ilia A. Solov’yov, Izabella Brand

## Abstract

Gram-negative bacteria possess a complex structural cell envelope that constitutes a barrier for antimicrobial peptides which neutralize the microbes by disrupting their cell membranes. Computational and experimental approaches were used to study a model outer membrane interaction with an antimicrobial peptide, melittin. The investigated membrane included di[3-deoxy-D-manno-octulosonyl]-lipid A (KLA) in the outer leaflet and 1-palmitoyl-2-oleoyl-sn-glycero-3-phosphoethanolamine (POPE) in the inner leaflet. Molecular dynamics simulations revealed, that the positively charged helical C-terminus of melittin anchors rapidly into the hydrophilic head-group region of KLA, while the flexible N-terminus makes contacts with the phosphate groups of KLA moving melittin into the boundary between the hydrophilic and hydrophobic regions of the lipids. Electrochemical techniques confirmed binding of melittin to the model membrane. To probe the peptide conformation and orientation during interaction with the membrane, polarization modulation infrared reflection absorption spectroscopy was used. The measurements revealed conformational changes in the peptide accompanied by reorientation and translocation of the peptide at the membrane surface. The study suggests that melittin insertion into the outer membrane affects its permeability and capacitance, but does not disturb the membrane’s integrity, indicating a distinct mechanism of the peptide action on the outer membrane of Gram-negative bacteria.

## Introduction

Increasing prevalence of Gram-negative bacteria to antibiotics imposes urgency on research for alternative ways of fighting bacterial infections.^1-4^Antimicrobial peptides (AMPs) belong to the innate immune system of animals and plants and are promising candidates as new antibiotic therapeutics. These peptides have a broad spectrum of activity against bacteria, viruses, fungi, parasites as well as cancer cells. In addition, the development of antimicrobial resistance against AMPs is less probable than against conventional antibiotics. Therefore, naturally occurring AMPs, their mutants and synthetic peptides are intensively screened as potential therapeutics.^1, 4, 5^AMPs neutralize microbes using different mechanisms of action. Most AMP interact with the bacterial cell envelope increasing the membrane permeability that leads to the cytoplasm leakage and cell death. However, some AMPs cross the membrane and enter the cell. The intracellular activity of AMPs involves their binding to DNA, inhibiting the protein synthesis machinery.^6-8^ AMPs are short, positively charged peptides that are characterized by large conformational flexibility, that usually have an undefined secondary structure in polar solutions. Electrostatic interactions constitute the driving force for anchoring of the AMPs on the membrane surface.^4, 7, 8^ Membrane associated AMPs interact with the lipid molecules, inserting themselves into the hydrophobic membrane fragment, while undergoing a conformational transformation stabilizing α-helical, *β*-sheet, or extended secondary structure motifs.^9-13^ Inserted AMPs disrupt the phospholipid membrane through the formation of pores, channels or lipid-peptide micelles.^13-16^ Different mechanisms of the membrane lysis by AMPs have been identified.^4, 9^

Melittin is the main component of the venom of western honey bee (*Apis mellifera*).^17^ This 26 amino acids long AMP is an amphiphilic molecule with a hydrophobic N-terminus and a cationic, polar C-terminus. In aqueous solutions, melittin monomers adopt a random coil conformation, while in solutions of high ionic strength, or peptide concentration, melittin aggregates into α-helical tetramers.^18^ Binding of melittin to a lipid bilayer is coupled with its folding into an α-helix. The peptide includes a proline residue at position 14, which is responsible for a bend between two helical fragments.^17, 18^ The presence of proline in the peptide sequence seems to enhance the antimicrobial activity of AMPs.^2^ Melittin became a model AMP for design of synthetic peptides having a broad cytolytic spectrum,^2, 3, 5^ and is also a common peptide for biophysical studies of AMPs interaction with models of biological cell membranes.^19-25^

The mechanism of the membrane disruption by melittin depends on two main factors: the peptide concentration and the lipid composition of the membrane.^14, 19, 26, 27^ In the initial stages of the interaction, melittin accumulates on the membrane surface. After reaching a certain coverage of the membrane surface, the hydrophobic N-terminus of melittin inserts into the hydrocarbon chain region of a lipid bilayer forming defects and pores.^14, 22, 23^ Several mechanisms of the phospholipid membrane lysis by melittin have been described, ^3, 9, 14^ however, the mechanism of neutralization of the cell envelope of Gram-negative bacteria is still not completely understood.

Gram-negative bacteria possess two membranes, the outer membrane (OM) and the inner membrane which are separated from each other by ca. 7 nm thick peptidoglycan layer.^28^ The OM is asymmetric and has an exceptional lipid composition; while the inner leaflet contains phospholipids. the OM leaflet is built of lipopolysaccharides, where Lipid A forms the amphiphilic part of each lipopolysaccharide. The polar head group of lipopolysaccharides is composed of saccharides that are subdivided into an inner core, outer core, and eventually O-antigen fragments. The carboxylic, phosphate and hydroxyl groups in the polar part of the lipopolysaccharides bind electrostatically divalent cations (Mg^2+^, Ca^2+^) to yield a rigid outer leaflet of the OM.^29-31^ Due to the structural and compositional complexity of the bacterial cell envelope the experimental characterization of its structure, physiochemical properties and functions are challenging. Computational approaches provide an emerging alternative to gain further insights on the structure, permeability, lipid mobility, and hydration of lipopolysaccharides in the OM.^32-39^

Molecular dynamics simulations and experimental results indicate that the action of AMPs on the OM is different as compared to the phospholipid bilayers.^7, 15, 34, 38-43^ Sharma *et al*. demonstrated for a random coil CM15 peptide approaching the external polar saccharide O-antigen and core regions of lipopolysaccharides preserves the peptide structure characteristic for aqueous solutions.^38^ The peptide penetrates the interfacial region of the inner core forming hydrogen bonds to phosphate groups of the lipid A. It was furthermore revealed that the CM15 peptide adopts a helical conformation upon entering the hydrophobic part of the membrane, which was also observed in another study for the LL-37 peptide interacting with the OM.^41^ The computational study of the CM15 peptide concluded that it has an ability to pass through the OM without disrupting it;^38^ it was hypothesized that CM15 peptide by lysing the inner phospholipid membrane neutralizes a microbe. Agglutination mechanism was proposed as another possible pathway of AMPs action on the bacterial membranes.^42^ Electrostatic binding of AMPs to lipopolysaccharides reduces the peptide mobility leading to its accumulation in the OM region, which by agglutination eventually induces bacterial cell death. The lack of clarity on the AMPs action on lipopolysaccharides in the OM calls for an in-depth analysis of the underlying molecular interactions.

In the present investigation we have studied melittin interaction with a model OM consisting of di[3-deoxy-D-manno-octulosonyl]-lipid A (KLA) in the outer leaflet and 1-palmitoyl-2-oleoyl-sn-glycero-3-phosphoethanolamine (POPE) in the inner leaflet as illustrated in Fig. 1. Molecular dynamics (MD) simulations were employed to describe the OM interaction with melittin at the atomic level, and to characterize the initial phases of melittin binding to the KLA-POPE model OM. The electrostatic interactions anchored the polar, positively charged C-terminus in the polar region of the KLA leaflet while the N-terminus displayed conformational and motional flexibility. The characteristic peptide conformation and orientation with respect to the OM surface have been determined. Long-time effects of the melittin binding to the KLA-POPE model OM were studied experimentally using *in situ* polarization modulation infrared reflection absorption spectroscopy (PM IRRAS) with electrochemical control. The results reveal that the electrostatic interactions between melittin and KLA affected the membrane stability and permeability. The association of melittin to the polar parts of KLA involves the peptide re-folding from a random coil to an α-helix. Finally, the measurements demonstrated a reorientation of melittin upon its association to the OM; over time melittin moves from the polar head groups in KLA to the hydrophobic core of the membrane changing the helix orientation from a tilted to normal to the membrane surface.

**Figure 1.**
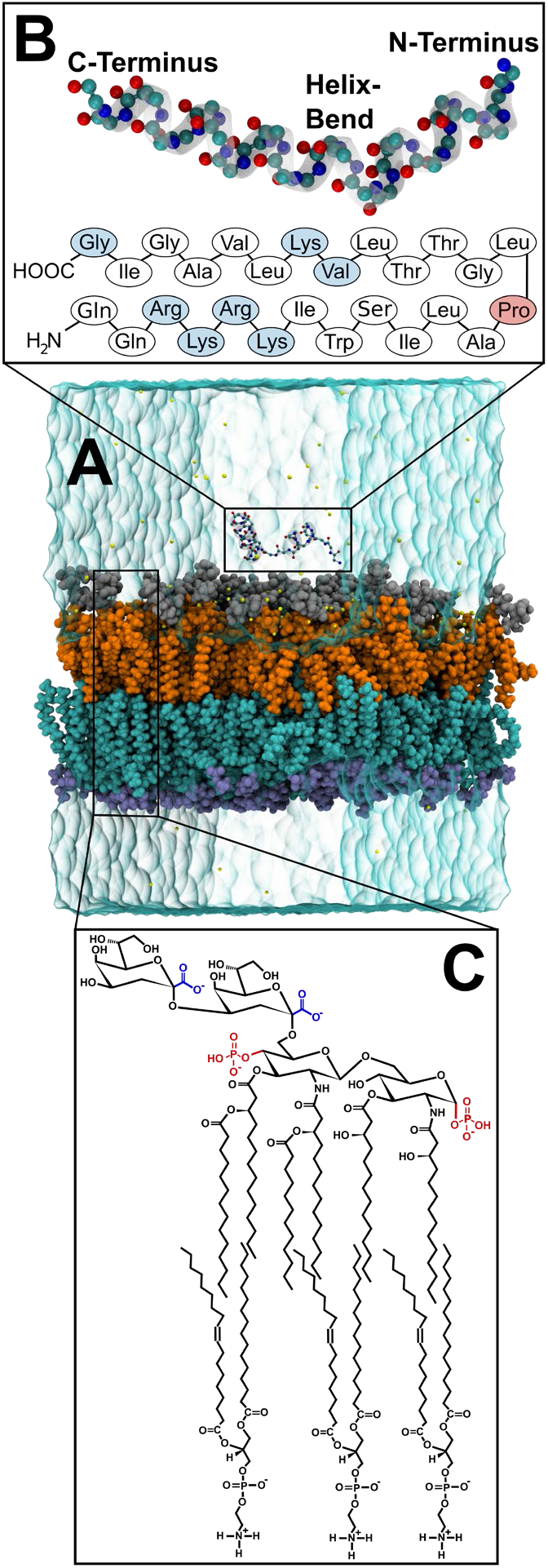
**A:** Molecular rendering of the melittin in a solvated membrane system. The carbon, oxygen and nitrogen atoms of the peptide are shown in cyan, red and blue, respectively. The different layers of the membrane, from bottom to top are the hydrophilic heads (purple) and the acyl chains (cyan) in POPE phospholipids, the lipid A (orange) and the inner core saccharides (silver) in KLA. Ions of various types are displayed as yellow spheres. **B:** Enlarged view of the peptide melittin and its primary sequence. The amino acids with the maximum contribution to the interaction energy are shown in light blue, Proline (Pro), where the helix bend in melittin occurs, is highlighted in red. **C:** The chemical structure of the membrane lipids KLA (top) and POPE (bottom). The phosphate groups and carboxylate groups in KLA are highlighted in red and blue,

## Methods

### Chemicals

1-palmitoyl-2-oleoyl-sn-glycero-3-phosphoethanolamine (POPE), *d*_31_-1-palmitoyl-2-oleoyl-sn-glycero-3-phosphoethanolamine (*d*_31_-POPE) and di [3-deoxy-D-manno-octulosonyl]-lipid A (ammonium salt) Kdo_2_-lipid A (KLA) were purchased from Avanti Polar lipids (USA). Lipids were used as received, no purification was done. Synthetic melittin (Cat. No.: M4171), Tris-(hydroxy-d-methyl)-amino-d_2_-methan (*d*_5_-TRIS), KClO_4_ (99.99%), Mg (ClO_4_)_2_×6H_2_O (99%), acid-sodium salt dihydrate (Na_2_EDTA) were purchased from Sigma Aldrich (Germany), NaCl (99.5%) and MgCl_2_ (>99% p.a.) were purchased from Carl Roth (Germany), Tris(hydroxymethyl)aminomethan (TRIS) from Fluka (Germany), ethanol and methanol from AnalaR Normapur, VWR (France), and D_2_O from Euroisotop (Germany).

### Langmuir-Blodgett transfer

Before each experiment, fresh lipid solutions were prepared. POPE was dissolved in CHCl_3_ while KLA was dissolved in CHCl_3_: CH_3_OH: H_2_O in 13:6:1 volume ratio. The concentration of POPE equalled 1 μmol ml^−1^ and the concentration of KLA was 1 mg ml^−1^. A microsyringe (Hamilton, USA) was used to place several μLs of the lipid solution at the liquid|air interface of the Langmuir trough (KSV Ltd., Finland). All aqueous solutions were prepared from ultrapure water [resistivity 18.2 MΩ cm (PureLab Classic, Elga LabWater, Germany)]. Before compression, the lipid solution was left for 10 min for the solvent evaporation. Surface pressure vs area per molecule isotherms were recorded using the KSV LB mini trough (KSV Ltd., Finland) equipped with two hydrophilic barriers. Surface pressure was recorded as a function of the mean molecular area. The accuracy of these measurements was ± 0.02 nm^2^ for the mean molecular area and ±0.1 mN m^-1^ for the surface pressure. Langmuir-Blodgett and Langmuir-Schaefer (LB-LS) transfers were used to prepare asymmetric supported planar KLA-POPE bilayers containing POPE in the inner (Au electrode oriented) and KLA in the outer (solution-oriented) leaflet on the gold surface, see Fig. S1. Prior to the LB-LS transfer each monolayer was compressed to the surface pressure of 30 mN m^−1^. First, the POPE monolayer was transferred from the aqueous subphase by a vertical LB withdrawing at a rate of 15 mm min^-1^. The transfer ratio was 1.10 ± 0.10. The POPE monolayer transferred onto the Au substrate was left for 3 hours. Next, a monolayer of KLA on 0.1 M KClO_4_ and 5 mM Mg(ClO_4_)_2_ aqueous subphase was compressed to the surface pressure of 30 mN m^−1^ and a horizontal LS transfer was used to fabricate the second leaflet on the gold surface, see Fig. S1. Planar lipid bilayers were dried for at least 2 hours before use in electrochemical and spectroelectrochemical experiments.

### Interaction of melittin with lipid bilayers

The concentration of melittin in a buffer solution [20 mM TRIS, 150 mM NaCl and 5 mM EDTA (pH=7.3 ± 0.1)] was set to equal either 1 or 10 μM. In such prepared solutions melittin exists in the monomeric form.^44^ Gold electrode modified with a lipid bilayer was incubated in a hanging meniscus configuration in the buffer solution containing the monomeric form of melittin. For the melittin concentration of 1 μM the incubation times were set to 15 min as well as 1 h. The incubation time from the 10 μM melittin solution was set to 15 min. After this time, the modified electrodes were carefully rinsed with water and used in either electrochemical or PM IRRAS experiments.

### Electrochemistry

Electrochemical measurements were performed in a glass three-electrode cell using a disc Au(111) single crystal (diam. 3 mm, MaTecK, Germany) as the working electrode in a hanging meniscus configuration. The surface roughness of the Au electrode was below 0.01 μm per 1 cm^2^ of the electrode surface. A gold wire was used as a counter electrode and an Ag|AgCl|sat.KCl (Ag|AgCl) as a reference electrode. All potentials are referred against the Ag|AgCl|sat.KCl electrode. The electrolyte solution was 100 mM KClO_4_ with 5 mM Mg(ClO_4_)_2_. A Methrom Autolab potentiostat (Methrom Autolab, Holland) was used to perform the electrochemical measurements. Prior to the experiment, the cell was purged with argon for 1 hour. The cleanness of the electrochemical cell was tested by recording the cyclic voltammograms in the electrolyte solution. Alternative current voltammetry (ACV) was used to measure the capacitance of unmodified and KLA-POPE bilayer modified Au(111) electrodes. AC voltammograms were recorded in negative and positive going potential scans at a rate of 5 mV s^-1^ and the perturbation of the AC signal of 20 Hz and 10 mV amplitude. The differential capacitance vs potential curves were calculated from the in-phase and out-of-phase components of the AC signal assuming the cell was equivalent to a resistor in series with a capacitor.

### Polarization modulation infrared reflection absorption spectroscopy

PM IRRA spectra were recorded using a Vertex 70 spectrometer with a photoelastic modulator (*f* = 50 kHz; PMA 50, Bruker, Germany) and demodulator (Hinds Instruments, USA). A home-made thin electrolyte layer spectroelectrochemical glass cell was washed in water, ethanol and placed in an oven (at 60 °C) for drying. CaF_2_ prism (optical window) was rinsed with water, ethanol and placed in UV ozone chamber (Bioforce Nanosciences, USA) for 10 minutes. The spectroelectrochemical cell has a build-in platinum counter electrode. The reference electrode was Ag|AgCl in 3M KCl either in D_2_O or H_2_O. A disc Au(111) single crystal (dia. 15 mm, MaTecK, Germany) was used as the working electrode and mirror for the IR radiation. The surface roughness of the Au electrode was below 0.01 μm per 1 cm^2^ of the electrode surface. A lipid bilayer was transferred on the working electrode surface using the LB-LS transfer. The electrolyte solution was 50 mM KClO_4_ with 5mM Mg(ClO_4_)_2_ in D_2_O. The electrolyte solution was purged for 1 hour with argon to remove oxygen. At each potential applied to the Au electrode 400 spectra with a resolution of 4 cm^-1^ were measured. Five negative and five positive going potential scans were recorded in each experiment. The negative going potential scan had the following potentials applied to the Au(111) electrode: 0.40, 0.25, 0.10, 0.00, –0.10, –0.20, –0.30, –0.40, –0.55, –0.65, –0.80 V while in the positive going potential scan: –0.80, –0.60, – 0.40, –0.30, –0.20, –0.10, 0.00, 0.10, 0.25, 0.40 V. At each potential the average, over five potential scans, spectrum was calculated and background corrected. The thickness of the electrolyte layer between the Au(111) electrode and the prism varied between 3 and 5 μm in different experiments. In one set of experiments the half wave retardation was set to 1600 cm^-1^ for the analysis of the amide I’ mode in melittin and C=O stretching mode in lipids. The angle of incidence of the incoming IR radiation was set to 55°. The solvent was D_2_O. For the analysis of the CD stretching modes in *d*_31_-POPE of the model OM, the the half wave retardation was set to 2100 cm^−1^. The angle of incidence was 52°. The solvent was H_2_O. All the spectra were processed using the OPUS v5.5 software (Bruker, Germany).

### Attenuated total reflection infrared spectroscopy

The analyte solution contained small unilamellar vesicles of the studied lipid mixture (KLA:POPE) (1.0: 3.2 mole ratio) and 4.4 × 10^*−*4^ mol L^*−*1^ melittin. 300 μl of 3.33 mg mL^*−*1^ POPE in chloroform and 300 μL of 3.33 mg mL^*−*1^ KLA in chloroform: methanol (2:1 vol) solutions were mixed and dried in a flow of argon. To remove the remaining solvent, the vials were placed in a vacuum desiccator for 72 hours. Next, 500 μL of 20 mM *d*_5_-TRIS, 150 mM NaCl and 5 mM MgCl_2_ in D_2_O was added to the dry lipid and the mixture was sonicated (EMAG-Technologies, Germany) at 35*°*C for 1 hour. Afterwards 300 μL synthetic melittin (Cat. No.: M4171) in 20 mM *d*_5_-TRIS, 150 mM NaCl and 5 mM MgCl_2_ in D_2_O was mixed with the vesicles solution and left for interaction for 60 min at 35 *°*C giving the analyte solution. Attenuated total reflection infrared spectra were recorded by co-adding 128 scans with a resolution of 4 cm^*−* 1^on a diamond prism using a MVP-Pro ATR unit (Harrick Scientific Products, Inc, USA) and the Bruker Vertex 70 spectrometer. The background spectrum was recorded for a drop (20 μl) of the electrolyte solution without lipid vesicles and melittin placed on the prism. The analyte spectrum was measured for a drop (20 μl) of the analyte solution (electrolyte with vesicles and melittin) placed on the prism. Subtraction of the analyte from the background spectrum gave the spectrum of lipids and melittin.

### Software used for MD simulations

MD simulations were carried out using NAMD 2.13 ^45, 46^ with the CHARMM36 force field ^47-49^ utilizing explicit water solvent in a TIP3P model ^50^. The computational platform VIKING ^51^ was used to set-up the computations. The membrane system was designed using CHARMM-GUIs ^52-54^ bilayer membrane builder ^53, 55-57^. Further preparation of the simulations and extraction of quantitative results, as well as the secondary structure analysis with STRIDE ^45^ were done using VMD^58^.

### MD simulations of solvated melittin

The structure of melittin monomer in its crystalline state was obtained from the melittin dimer available in the protein databank ^59, 60^ (PDB), with the ID 2mlt entry.^61^ In the crystalline state melittin forms two α helices which are separated by a bend occurring at the 14th residue, see Fig. 1B. For the equilibration simulation, the melittin monomer was solvated in a cubical water box with a side length of 9 nm. The salt concentration used in the simulation resembled the experimental value, of 50 mM KCl and 5 mM MgCl_2_, resulting in a total of 69585 atoms in the system. The system was simulated for 100 ns, imposing periodic boundary conditions and a constant temperature of 303.15 K controlled by a Langevin thermostat.^62^ The simulation was split into 3 simulation stages. In the first stage, a 10 ns simulation was performed, employing the isothermal-isobaric statistical ensemble (NPT), maintaining a pressure of 1 bar using a Nosé-Hoover-Langevin piston pressure control ^62, 63^ with a timestep of 1 fs for integration of the equations of motion. The second and the third stages used canonical ensemble (NVT) with an integration timestep of 1 fs and 2 fs and included simulations of 5 ns and 85 ns duration, respectively. No constraints were imposed on the atoms of the system during the three simulation stages (equilibration simulations). Explicit non-bonded interaction were neglected at 12 Å with a set smooth switching distance starting at 10 Å. Electrostatic interactions beyond 12 Å were calculated using the particle-mesh Ewald method.^64^ After the equilibration simulations the system was further simulated for another 200 ns for data acquisition (production simulation). The acquired data was used as an independent control simulation of melittin in water.

### Equilibration of the bilayer membrane

The bilayer membrane was designed using CHARMM-GUIs bilayer membrane builder.^52-54^ The chemical structure of the simulated OM matches the membrane studied in the experiments and is shown in Fig. 1C. The negative charge at phosphate groups in the lipid A part in KLA were neutralized using Mg^2+^ ions while the inner core in KLA using Na^+^ ions. The membrane had the density of 96.37 atoms nm^*−*3^. The number of lipids in the model OM are 34 KLA in the outer leaflet and 109 POPE lipids in the inner leaflet. Before the equilibration procedure KLA and POPE molecule occupied an area of 1.95 nm^2^ and 0.61 nm^2^, respectively (see Section S2 and Fig. S2); the converged equilibrated values appear 1.87 nm^2^ and 0.58 nm^2^, respectively. These values agree well with the LB-LS transfer conditions at which the average area per lipid *A*_*KLA*_ was 1.96 nm^2^ and *A*_*POPE*_ - 0.67 nm^2^.^65^ The membrane was solvated in a water box, matching the membrane area of (8 × 8) nm^2^ and having a height of 11 nm. The concentration of KCl was assumed 50 mM. Due to the low amount of solvent the 5mM MgCl_2_ used in the experiments, the Mg^2+^ ions were omitted in the simulation. NAMD^66^ was used to equilibrate the membrane for a total of 200 ns using an isothermal-isobaric ensemble and periodic boundary conditions in 3 distinct stages. The CHARMM36 force field^48, 49, 67^ was employed in the simulations. A constant temperature of 303.15 K controlled by a Langevin thermostat^62^ and a constant pressure of 1 bar using a Nosé-Hoover-Langevin piston pressure control ^62, 63^ was imposed during the whole simulation. The first stage was simulated for 5.375 ns with an integration timestep of 1 fs. The second and the third stages used an integration timestep of 2 fs each and were simulated for 2.125 ns and 90 ns, respectively. The simulations in stage 1 and stage 2 considered the movement of the head groups of all POPE and KLA lipids constrained using a harmonic restraint potential. The force constant of that restraint was decreased from 5 to 0.2 kcal mol^-1^ during the simulation stages 1 and 2. Explicit non-bonded interactions with a smooth switching starting at 10 Å were cut off at 12 Å and electrostatic interactions were calculated using the particle-mesh Ewald method^64^ beyond the distance of 12 Å.

### Equilibrium molecular dynamics melittin atop bilayer membrane

A combined system, shown in Fig. 1A, of the equilibrated melittin monomer and the equilibrated membrane was used to generate data for analysis. The melittin monomer, which acquired a V-shape during the equilibration simulation was placed on top of the membrane in three different initial orientations, depicted in Fig 2. Simulation 1 (Sim. 1) considered melittin placed with its helix bend 1.2 nm above the membrane surface (Fig. 2A). Simulation 2 (Sim. 2, Fig. 2B) and simulation 3 (Sim. 3, Fig. 2C) had melittin placed with the C-terminus and the N-terminus nearest to the membrane, respectively. All systems were ionized with a salt concentration of 50 mM KCl, omitting the 5mM MgCl_2_ used in the experiments. These composite systems with a total number of atoms varying between 82399 and 82405 were simulated with periodic boundary conditions. After the composite systems were constructed, each system was first equilibrated for 5 ns in the isothermal-isobaric ensemble, maintaining a pressure of 1 bar using a Nosé-Hoover-Langevin piston pressure control ^62, 63^ and a constant temperature of 303.15 K using a Langevin thermostat ^62^, followed by a 5 ns simulation in the canonical ensemble with an integration timestep of 1 fs. After the equilibration simulation, production simulations of each system with a duration of 200 ns in the canonical ensemble with an integration timestep of 2 fs were performed. A 12 Å cutoff was imposed for the non-bonded interactions, which were linearly switched to zero starting from 10 Å. Beyond these distances, electrostatic interactions were calculated using the particle-mesh Ewald method.^64^

**Figure 2.**
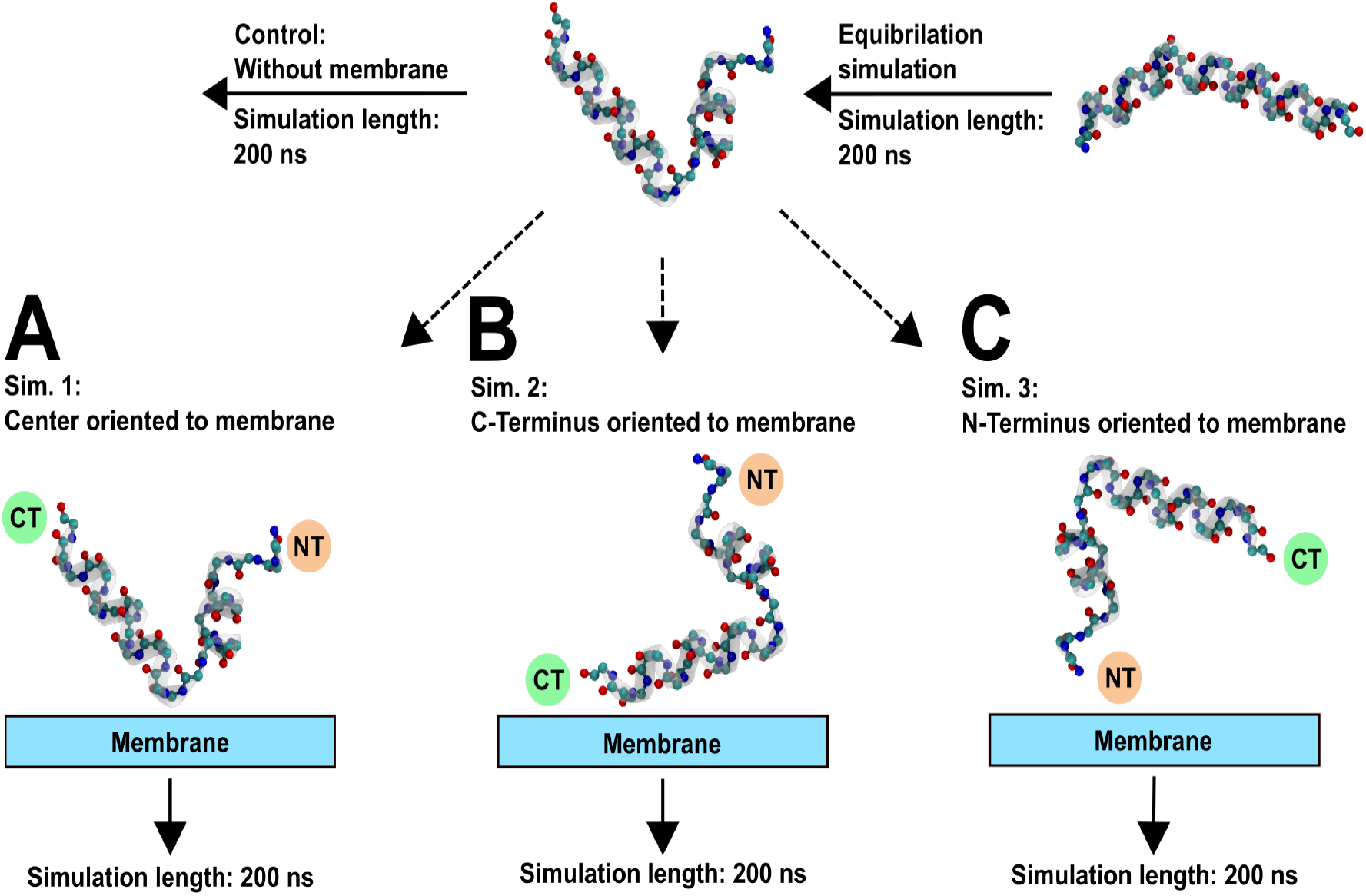
Schematic representation of the simulation procedure. The melittin peptide was equilibrated for 200 ns solvated in a cubic water box. The resulting peptide conformation was used for the four consecutive simulations. The peptide was placed on-top of the equilibrated membrane in three different orientations facing the membrane’s surface: **A**: with the bend in the peptide’s middle part, Sim. 1; **B**: with its C-terminus, Sim. 2 and C: N-terminus, Sim. 3. An independent control simulation of melittin without a membrane solvated in a cubic water box was also carried out (upper panel). Water is omitted in the graphical representations for clarity.

## Results and Discussion

### Electrochemical Characterization of the Model Outer Membrane Interacting with Melittin

Electrochemical measurements provide information about the compactness and stability of model lipid membranes in changing electric fields. The capacitance of the Au electrode depends on the chemical nature of the adsorbed species, their surface coverage, and packing. The capacitance changes as a function of the applied potential. The difference between the applied potential (*E*) and the potential of zero charge (*E*_pzc_) is an adequate approximation of the membrane potential (*E*^m^),^65, 68^ which reads as

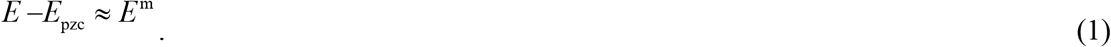

The membrane potential has an important biological significance and indicates directly changes in cell membranes.^69^ Figure 3 shows the capacitance of the KLA-POPE bilayers with and without the bound melittin. Exposure of the OM to melittin changed the electrochemical properties of the membrane in a peptide concentration-dependent manner. In one set of experiments, the OM was immersed for 15 min into a solution containing either 1μM or 10 μM melittin. Afterwards, these membranes were transferred to the electrolyte solution containing 50 mM KClO_4_ and 5 mM Mg(ClO_4_)_2_. An increase in the melittin concentration in the electrolyte solution leads to an increase in the membrane capacitance (Fig. 3, lines a-c). A broad capacitance maximum appears at *−*0.30 V < *E*^m^ < 0.05 V, indicating a phase transition and/or a molecular scale rearrangement in the bilayer. In the OM with melittin associated from the solution phase, the membrane potential of this peak shifts towards more negative values compared to the pure KLA-POPE bilayer (Fig. 3, line a). At *E*^m^ <*−*0.8 V, the measured capacitance was negligibly affected by the presence of melittin (Fig. 3, lines a-c). This result means that the presence of melittin has no effect on the potential-driven desorption and disruption of the OM. The interaction of melittin with the OM affected the membrane permeability and stability, but it did not lead to dissolution of the lipid bilayer from the Au(111) surface.

**Figure 3.**
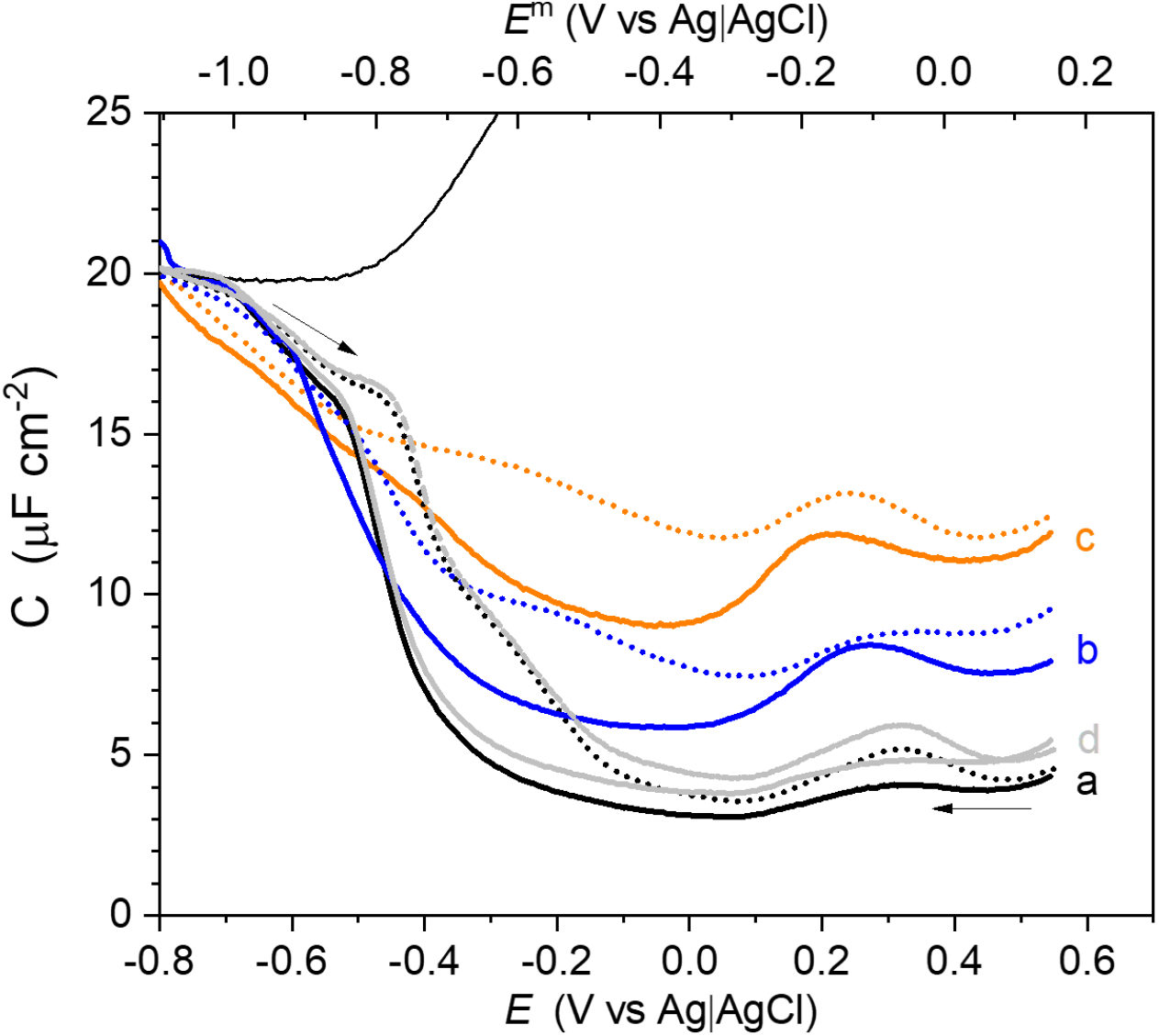
Capacitance (*C*) versus potential (*E*) and membrane potential (*E*^m^) of the KLA-POPE (a-c) bilayers on the Au(111) electrode surface. The measurements were done in the absence of melittin (a, black); after 15 minutes of interaction with 1 μM melittin (b, blue); and 15 minutes of interaction with 10 μM melittin (c, orange). Capacitance for specially prepared LB-LS KLA:Mel (9:1 mole ratio)-POPE bilayer is shown (d, gray). Solid and dotted lines correspond to the negative and positive going potential scans, respectively. Arrows show the directions of the potential scans. Thin black line: Capacitance of the unmodified Au(111) electrode. 50 mM KClO_4_ and 5 mM Mg(ClO_4_)_2_ was used as the electrolyte solution.

In the second set of experiments, a KLA : melittin (9:1 mole ratio) monolayer was transferred by the LS method onto the Au(111) surface covered by the inner POPE leaflet. In this case the electrochemical characterization of the lipid-peptide membrane was comparable to that of the pure KLA-POPE bilayer (Fig.3, lines a and d), indicating an insignificant effect of melittin on the electrochemical properties of the OM. Thus, the interaction pathway of melittin with the OM depends on the delivery strategy of the peptide.

Melittin may either accumulate on the surface of the KLA leaflet, or penetrate into the OM forming defects and channels as schematically shown in Fig. S3. Electrochemical measurements are sensitive to different supramolecular arrangements of molecules in a film covering an electrode surface. The dielectric constant of the adsorbing molecule has an impact on the capacitance value, see Section S3. Lipid molecules are amphiphilic and contain long acyl chains. The dielectric constant of a hydrocarbon chain is *∼*2 while that of the polar head group of lipids is 8 - 10 ^70^; the dielectric constant of proteins is difficult to estimate and was reported to be in the range 5 - 30^71^. Experimentally measured capacitance of the OM-melittin films contains a contribution from the capacitance of the lipid membrane (*C*_m_), adsorbed peptide (*C*_p_) and the diffuse layer (C_*dl*_), and reads as:

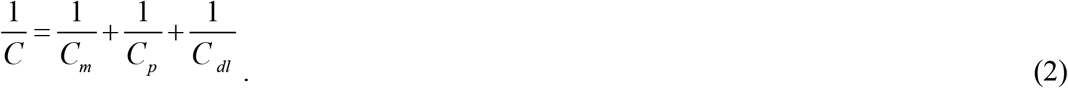

Adsorption of melittin on top of the lipid bilayer surface, see Fig. S3A, does not affect the measured capacitance, because *C* is dominated by the capacitance of the subsystem with the lowest value (*C*_m_). In comparison to a pure KLA-POPE bilayer (see Fig. 3, line a), the OM with KLA and melittin transferred by the LB method (see Fig. 3, line d) display minor changes in the capacitance. The result suggests that melittin accumulates in the polar head group region (on top) of the KLA monolayer and does not enter the hydrophobic fragment of the KLA lipid. In contrast, an increase in the capacitance observed upon melittin binding from the solution, compared to the pure KLA-POPE bilayer, indicated an insertion of the peptide into the membrane, see Fig. S3 B,C. The membrane insertion of melittin is expected to affect the packing and orientation of the lipid molecules and change the conformation/hydration of the peptide in the environment of the OM. The electrochemical data suggest molecular-scale rearrangements in the orientation and conformation of lipids and melittin during interaction. To describe this lipid-peptide interaction at molecular and atomic levels MD simulation (short interaction time) and *in situ* spectroelectrochemical (long interaction time) experiments were performed.

### Melittin binding, conformation and orientation: A short-time scale of the interaction

Three independent simulations (Sim. 1 – 3, see Fig. 2) show melittin binding to the model membrane. To gain insight into the binding mechanism, the interaction energies between the individual amino acids in the peptide and the membrane were calculated. Interaction energies include Coulomb and van der Waals contributions. The details of the calculation of the electrostatic and van der Waals contributions to the interaction energy are described in the Supporting Information S4. Figure 4A shows the time evolution of the total interaction energy (*E*_B_) between the peptide and the membrane for all three simulations (Sim. 1 - 3). The figure features a decrease in the interaction energy in all simulations, which indicates binding of melittin to the KLA leaflet after 200 ns, see Fig. 4A. Note that the different initial orientations of the peptide with respect to the membrane did not significantly affect the resulting melittin binding energy, however affected how fast the stable value was reached. Comparing the results of Sim. 3 (purple) with Sim. 1 (orange) and Sim. 2 (lilac), it is clear that the peptide in Sim. 3 experienced an unfavorable initial orientation of the N-terminus above the membrane surface. In this case, during the first 100 ns of the simulation, the peptide moved away from the membrane, reoriented, and finally bound with its C-terminal to the membrane.

**Figure 4.**
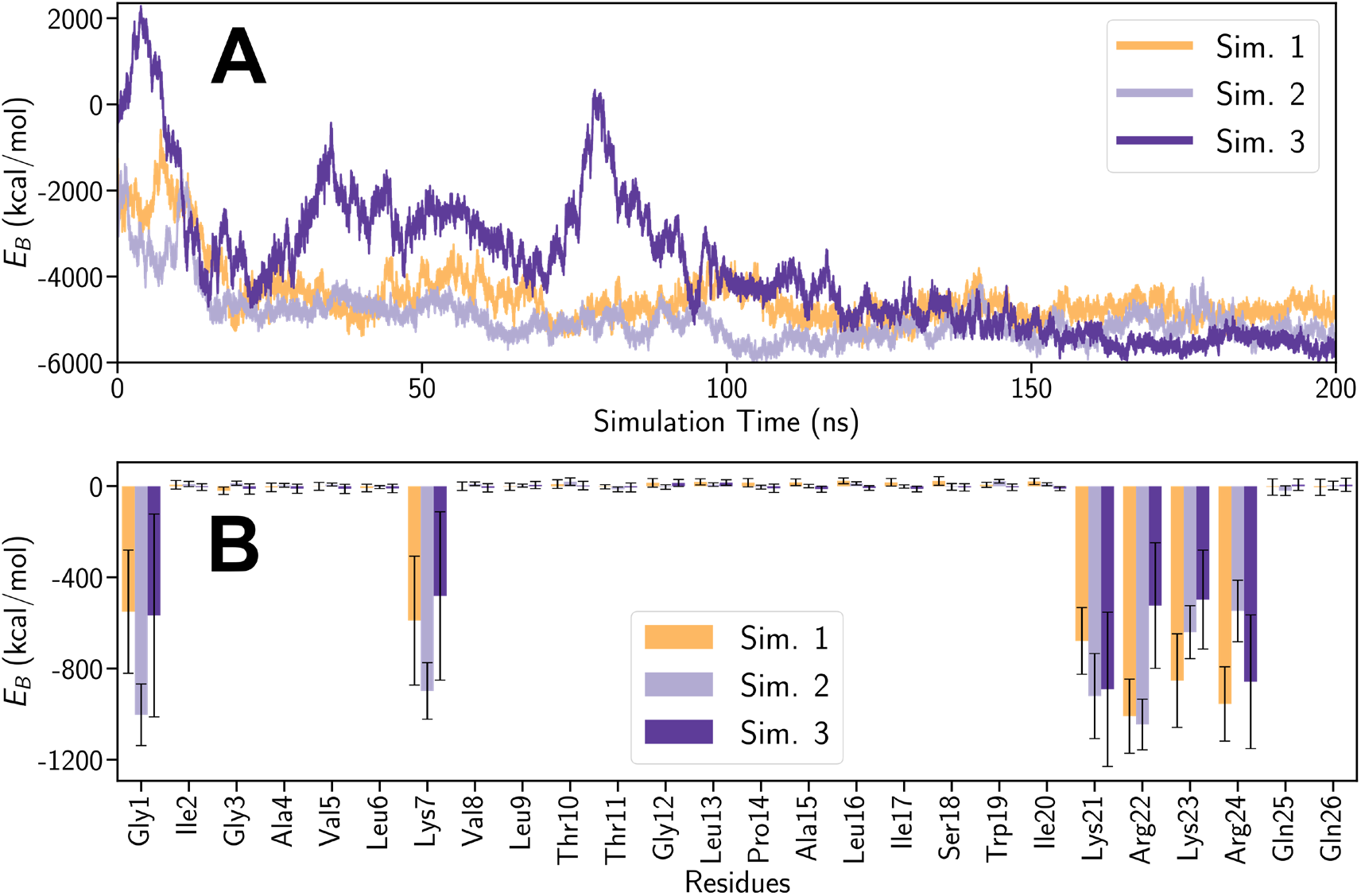
Interaction energy *E*_B_ between melittin and the KLA-POPE membrane calculated for the simulations Sim. 1, Sim. 2 and Sim. 3. **A:** Temporal evolution of *E*_B_. **B:** Time-averaged values of *E*_B_ computed for each individual amino acid residue in the peptide. The standard deviations are shown as error bars.

The individual contributions of each residue to the overall interaction energy are shown in Fig. 4B. From the 26 residues of melittin, only six contributed significantly to the overall interaction energy: Gly1, Lys7, Lys21, Arg22, Lys23 and Arg24. The positively charged Lys and Arg residues (21 - 24) present in the helix of the C-terminus bind the peptide to the OM. Hydrogen bonds between the residues Lys21, Arg 22, Lys23 and Arg24 of the peptide and the KLA in the OM were observed. The ε-ammonium group in Lys and the guanidinium group in Arg acted as the hydrogen donors, while the oxygen atoms of carboxylate, phosphate and hydroxy groups in KLA accepted the hydrogen atoms. The measured donor-acceptor distances were between 2.2 Å and 3.0 Å, depending on the specific groups that formed the hydrogen bond. With advancing simulation time these bonds were observed more frequently indicating an increase in the stability of the bonds and thereby explaining the strong binding of the helical structure at the C-terminus of melittin to the OM. At the N-terminus only Gly1 and Lys7 residues interacted with the membrane. Binding of melittin to the OM is connected with the stabilization of the α-helix secondary structure, see Fig. S4. The helices constituted 50 *−* 75% of the overall peptide structure. In contrast, in the control simulation performed in water, a loss of helicity to ca. 35% over 200 ns was observed, see Fig S4A.

Interestingly, the final configurations of melittin adsorbing on the KLA leaflet of the OM obtained in all three simulations, are very similar, see Fig. S5. The peptide binds with its positively charged helical C-terminus to the membrane. The secondary structure of the N-terminus of melittin was poorly defined and displayed some motional freedom. However, the Gly1 residue in the N-terminus of melittin made close contacts leading to the formation of hydrogen bonds to the negatively charged phosphate and carboxylate groups in KLA. The interaction of the Gly1 residue in melittin with the OM was analyzed in detail, by studying the formation of hydrogen bonds between the N-terminus and the phosphate group in the lipid A part and carboxylate groups of the inner core in KLA molecules. The results are illustrated in Fig. 5A.

**Figure 5.**
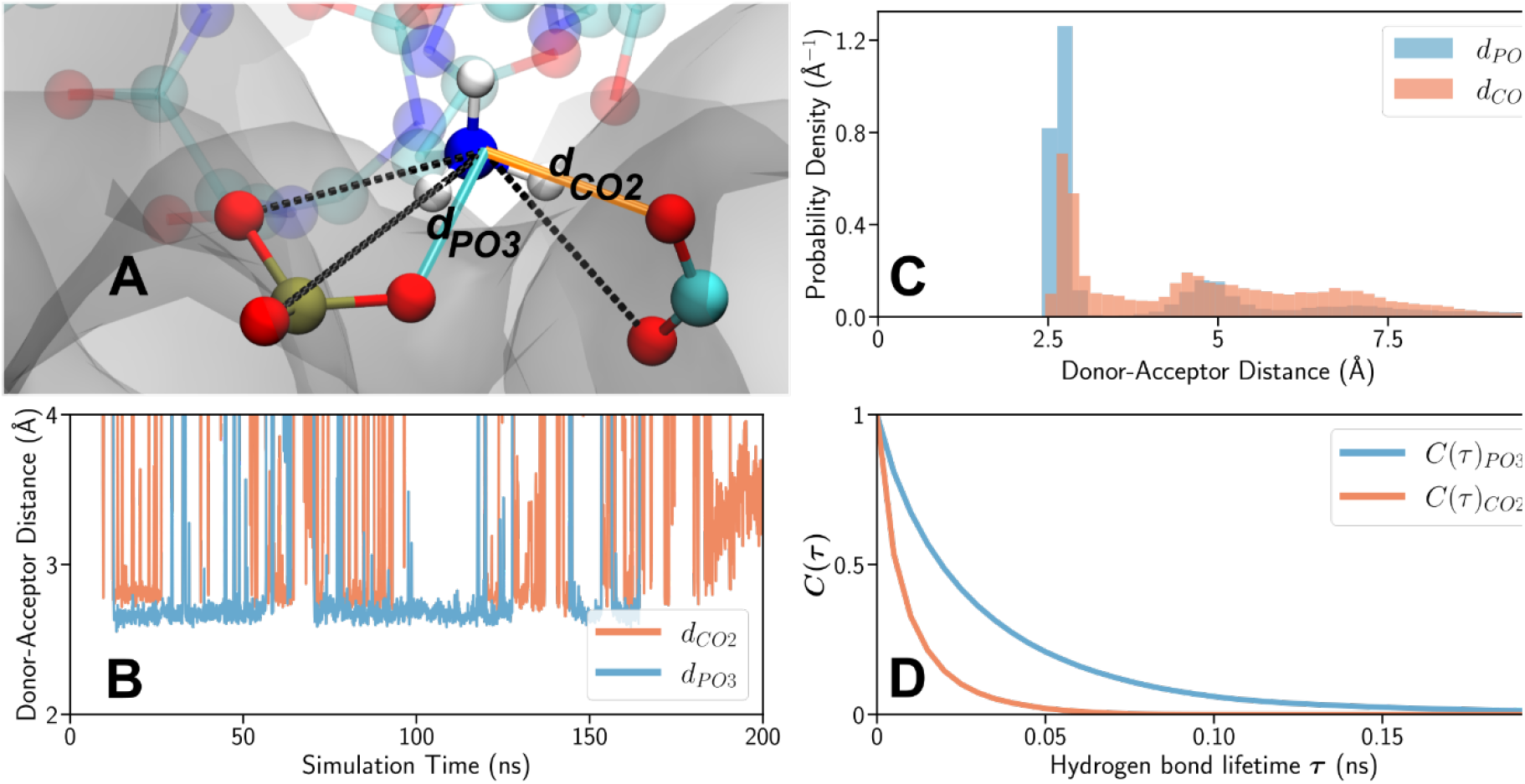
**A:** Rendering of a phosphate group in the lipid A part a carboxylate group in the inner core in KLA in close proximity to the N-terminus of melittin. The nitrogen, carbon, phosphorus, oxygen and hydrogen atoms are shown in blue, cyan, ochre, red and white, respectively. The membrane is visualized as a gray surface and the protein backbone atoms are shown in the corresponding translucent colors. The black dotted lines correspond to the distances between each oxygen atom of the carboxylate and phosphate groups and the geometric center of the three hydrogen atoms of the protein N-terminus. The minimal distances to the carboxylate group and to the phosphate group are labeled *d*_*PO3*_ and *d*_*CO2*_, respectively and are highlighted in orange and cyan. **B/C:** The minimal distance between the geometric center of the hydrogen atoms in the amine group in Gly1 in the N-terminus and the phosphate group *d*_*PO3*_ (cyan) and the carboxylate groups *d*_*CO2*_ (orange) calculated from Sim. 1 and plotted over simulation time (**B**) and shown as a histogram (**C**). **D**: The hydrogen bonding time autocorrelation function, see Eq. (3), of the phosphate group *C*_*PO3*_*(τ)* (cyan) and the carboxylate groups *C*_*CO2*_*(τ)* (orange).

A hydrogen bond is defined by geometric criteria that includes the distance between the donor and acceptor atoms and the value of the donor-hydrogen-acceptor planar angle.^72^ The NH_2_ group in Gly1 in the N-terminus and a phosphate or carboxylate group were considered hydrogen-bonded if the donor-acceptor distance *d* was less than 3 Å and the value of the donor-hydrogen-acceptor angle turn out to be 150°-180°. Figure 5A shows the typical location of the amide group in Gly1 in the N-terminus of melittin (nitrogen in blue and hydrogen in white) in the KLA leaflet of the OM (gray surface). Gly1 interacts with the phosphate groups of the lipid A part in KLA (phosphorus in ochre, oxygen in red) and the carboxylate groups of the inner core in the KLA molecule (carbon in cyan, oxygen in red). Black dotted lines in Fig. 5A represent the existing donor-acceptor distances between oxygen atoms in the PO_3_ and 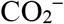 groups of KLA and the N-terminus of melittin. The orange line represents the minimal donor-acceptor distance to the closest oxygen atom of the 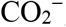group and the cyan line represents the minimal distance to the closest oxygen atom of the PO_3_ group. The minimal distances obtained from the Sim. 1, *d*_*CO2*_ and *d*_*PO3*_, are plotted over the simulation time in Fig. 5B as well as a probability distribution in Fig. 5C. The most probable donor-acceptor distance between the NH_3_ group of the Gly1 residue and the oxygen atoms of the phosphate groups is smaller, compared to the distance to the carboxylate group. The distance *d*_*PO3*_ is rather conserved over significant simulation time at a value of *∼*2.7 Å, while the value of *d*_*CO2*_ fluctuates more, indicating a stronger and longer living hydrogen bond to the phosphate groups. The distance between the donor-acceptor sites alone, however, is insufficient to define the lifetime of a hydrogen bond since it also depends on the planar angle formed by the involved groups. For further investigations of the hydrogen bond dynamics, the average lifetime (*T*) was calculated through the autocorrelation function defined as ^79^:

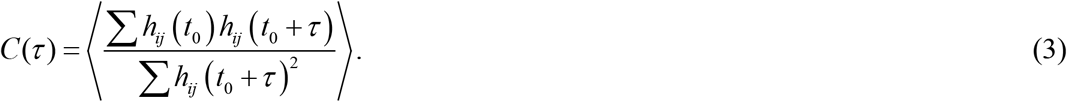

Here *h*_*ij*_(*t*_0_) measures weather a pairing between a hydrogen atom *i* of the N-terminus and the acceptor oxygen atom *j* of the carboxylate or phosphate groups satisfy the hydrogen bond criteria at the reference time instance *t*_*0*_, while *h*_*ij*_ (*t*_0_ *+τ*) checks for the hydrogen bond’s existence at the time instance *t*_0_ *+τ* . The summation was carried out over possible pairs *ij* between the hydrogen atoms of the NH_3_ group of Gly1 and the oxygen atoms of either the phosphate group or carboxylate group of KLA. Angular brackets indicate an average over the different starting time instances.

The computed autocorrelation functions are shown in Fig. 5D for the hydrogen bonds formed to the oxygen atoms of the phosphate groups *C*_*PO3*_*(τ)* and to the oxygen atoms of the carboxylate groups *C*_*CO2*_*(τ)*. The hydrogen bond lifetime *T* is then defined as the integral of the autocorrelation function^72^ as:

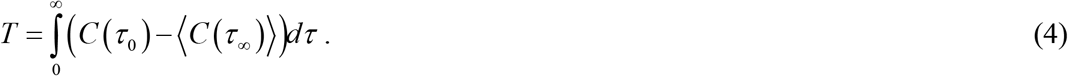

The lifetime *T* is calculated by fitting the results of the autocorrelation function with a bi-exponential function and subsequently numerically integrating Eq. (4). This yields an average lifetime of 0.03 ns for a hydrogen bond between the N-terminus in melittin and the oxygen atom of a phosphate group in KLA and 0.01 ns for a hydrogen bond with the carboxylate group in KLA. The presence of the charged N-terminus and its binding to the carboxylate and phosphate groups might supersede a divalent cation that binds to a phosphate group or a carboxylate group of the KLA in a melittin free membrane. The earlier PM IRRAS results indicated that upon melittin binding the coordination to the carboxylate groups in the inner core of the KLA changes, conforming that the negatively charged residues in KLA interact directly with melittin.^73^ MD simulations show also that the N-terminus in melittin, within 200 ns of the simulation time, penetrated the inner core to make stable hydrogen bonds with the phosphate residues located at the polar-hydrophobic interface of the lipid A part in the KLA. Binding of melittin to the KLA may change the balance of charges in the outer leaflet of the OM and cause structural disturbances in the lipid membrane, which is an essential part of the insertion of melittin into the membrane.^74^

The tilt angle of the helices in melittin relative to the membrane surface was determined. Each amide group of melittin includes an IR-active C=O bond aligning with the corresponding transition dipole moment. The dipole moment for each residue of melittin is characterized by a tilt angle *θ*_n_, computed relatively to the membrane surface normal as

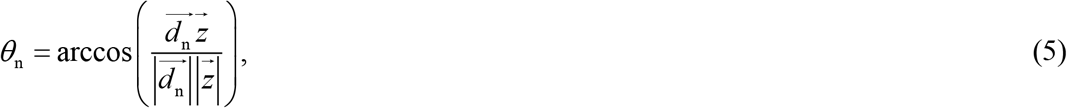

where 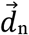 is the transition dipole moment of the C=O stretching mode (*ν*(C=O) mode) in the n-th residue and 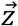 is the normal vector pointing perpendicular to the membrane surface. The resulting time average angle ⟨θ⟩ for the whole melittin can be computed as:

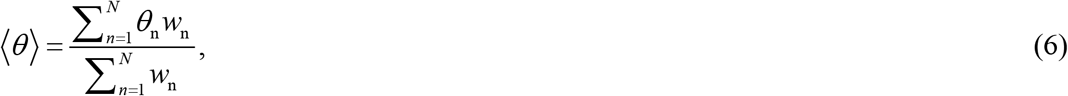

where the weights *w*_n_ describe the coupling of the transition dipole moments of the *ν*(C=O) modes in melittin to the electric field vector of the reflected IR radiation and are defined as the *z*-component of the normalized vector of the respective dipole moment 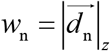. The summation was carried out over 600 ns of the combined trajectories of the simulations Sim. 1-3, yielding the average tilt angle of ⟨*θ*⟩ = 48.03°. The tilt of the long axis of the helix in melittin with respect to the membrane surface normal could readily be related to ⟨*θ*⟩ (see section S10), and ranges between 36 ° - 39 °.

### Melittin binding, conformation and orientation: Long interaction time scale

Figure 6 shows the PM IRRA spectra of the KLA-POPE bilayer after 15 minutes of incubation in 1 μM melittin solution. The spectra feature the *ν*(C=O) stretching modes in the ester carbonyl groups in lipids, amide I’ vibration mode mainly in melittin and *ν*_as_(COO^*−*^) stretching mode in KLA. Furthermore, two amide groups in KLA contribute also to the amide I’ vibration mode, however these groups have a slight impact on the resulting IR spectrum.

**Figure 6.**
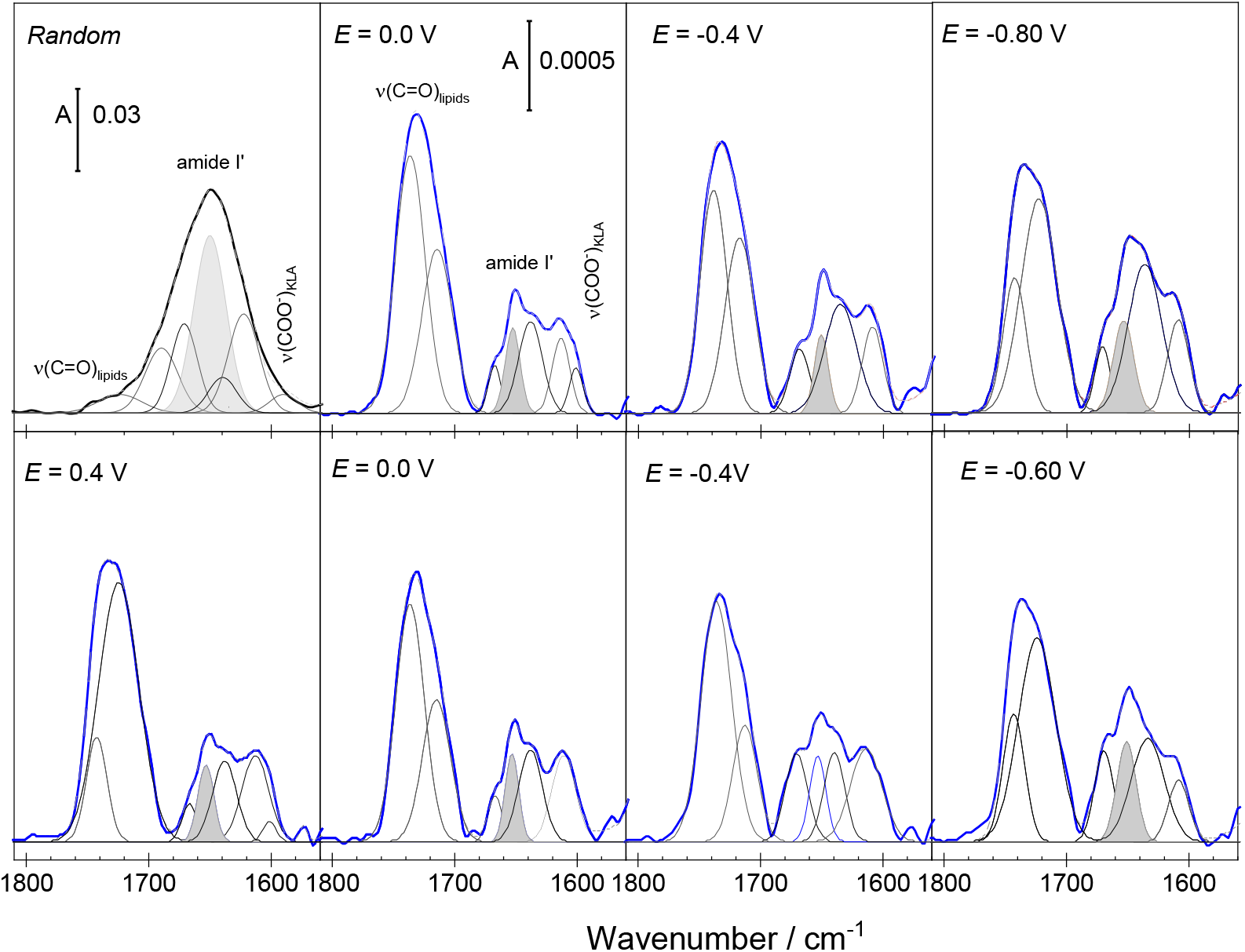
IR spectra of KLA-POPE bilayer systems (thick lines) after interaction with melittin. The left spectrum in the upper panel shows an ATR IR spectrum of KLA-POPE vesicles after 60 minutes interaction with 4.4 *×*10^*−*4^ M melittin. Other figures show the PM IRRA spectra of KLA-POPE bilayers after 15 minutes of interaction with melittin (blue lines) recorded at different electrode potentials. The upper and lower panels show results for the negative and positive scans, respectively. Thin black lines show the band deconvolution results. The bands highlighted in gray show the amide I’ mode of *α*-helices in melittin. Measurements were carried out for 1 μM melittin in 50 mM KClO_4_ and 5 mM Mg(ClO_4_)_2_ in D_2_O. The absorbance is shown in arbitrary units.

In the presence of melittin the *ν*_as_(COO^*−*^) absorption bands in KLA appear at 1610-1608 cm^*−*1^ and 1600 cm^-1^. The absorption maximum of the high wavenumber *ν*_as_(COO^*−*^) mode is ca. 5 cm^*−*1^ down-shifted compared to the position of this band in the pure KLA-POPE bilayer, ^65,73^ being the result of changes in the coordination to the carboxylate group. The observed spectral changes are associated to the removal of Mg^2+^ ions and formation of direct hydrogen bonds to positively charged amino acids in melittin.

The amide I’ band in melittin is a complex spectral feature centered at 1650 cm^*−*1^, see Fig. 6. The IR spectrum of melittin KLA-POPE vesicles corresponds to a random distribution of the peptide. The amide I’ band was deconvoluted into five components centered at 1689 cm^*−*1^, 1670 cm^*−*1^, 1650 cm^*−*1^, 1639 cm^*−*1^, and 1622 cm^*−*1^, see Fig. 6 and Fig. S6A. The complexity of the amide I’ mode points at structural flexibility and significant differences in the hydration and hydrogen bonding network in melittin in solution and lipid bilayer associated states, agreeing with earlier findings.^22, 23, 75-78^ For randomly distributed melittin, the strongest component of the amide I’ band (1650 cm^*−*1^) stems from α-helices that constitute 42% of the peptide secondary structure. The modes at 1670 cm^*−*1^ and 1630 cm^*−*1^ could be attributed to either helical (3_10_- or *π*-) structures or random coils.^77^ Assignment of the IR absorption modes at 1689 cm^*−*1^ and 1622 cm^*−*1^ is primarily associated with antiparallel *β*-sheets.^75, 78^ Drastic changes in the hydrogen bonding network at the helical and disordered peptide fragments, however, may result in similar spectral changes. Dehydration of the C=O group causes an up-shift of the amide I’ mode to 1685-1700 cm^*−*1^.^79^ A strong hydration of the helical peptide fragments may result in a down-shift of the amide I’ mode to ca. 1625 cm^*−*1^.^23, 77, 80^ The IR spectrum of melittin in solution was measured for the peptide:lipid ratio of 1:22. At such a high content of melittin, the peptide may aggregate on the vesicle surface that may induce even a conformational change to *β*-sheet structures as proposed earlier.^23, 75, 78^

MD simulations confirm the structural flexibility of non-bonded melittin. In the control simulation, in water, gradual loss of the helical structure to ca. 35% was observed, see Fig. S4A. Simultaneously, changes in the backbone dihedral angles indicate that melittin may adopt a *β*-sheet structure, see Fig. S4C. It is therefore reasonable to conclude that melittin approaching the OM surface from the solution phase is characterized by a conformational flexibility and hydrogen bonding network of different strengths.

After 15 minutes of the OM interaction with 1μM melittin, the amide I’ band features three components centered at 1669 cm^*−*1^, 1653 cm^*−*1^ and 1636 cm^*−*1^, see Fig. 6 and Fig. S6B. The observed spectral changes indicate that membrane associated melittin is composed of helices and random coils, reflecting only to a certain extent the structural flexibility found in the solvated peptide, see Fig. 6.

Figure 7 shows the PM IRRA spectra of the OM after 15 minutes interaction with 10 μM melittin. Independently of the electrode potential a strong symmetric amide I’ band, centered at 1648 cm^*−*1^, is present in the spectra. At *E* > *−*0.4 V the amide I’ band in melittin contains one component assigned to α-helices. Described above results of MD simulations also predict a stabilization of the helical structure of melittin in the membrane bound state, see Fig. S4A,B.

**Figure 7.**
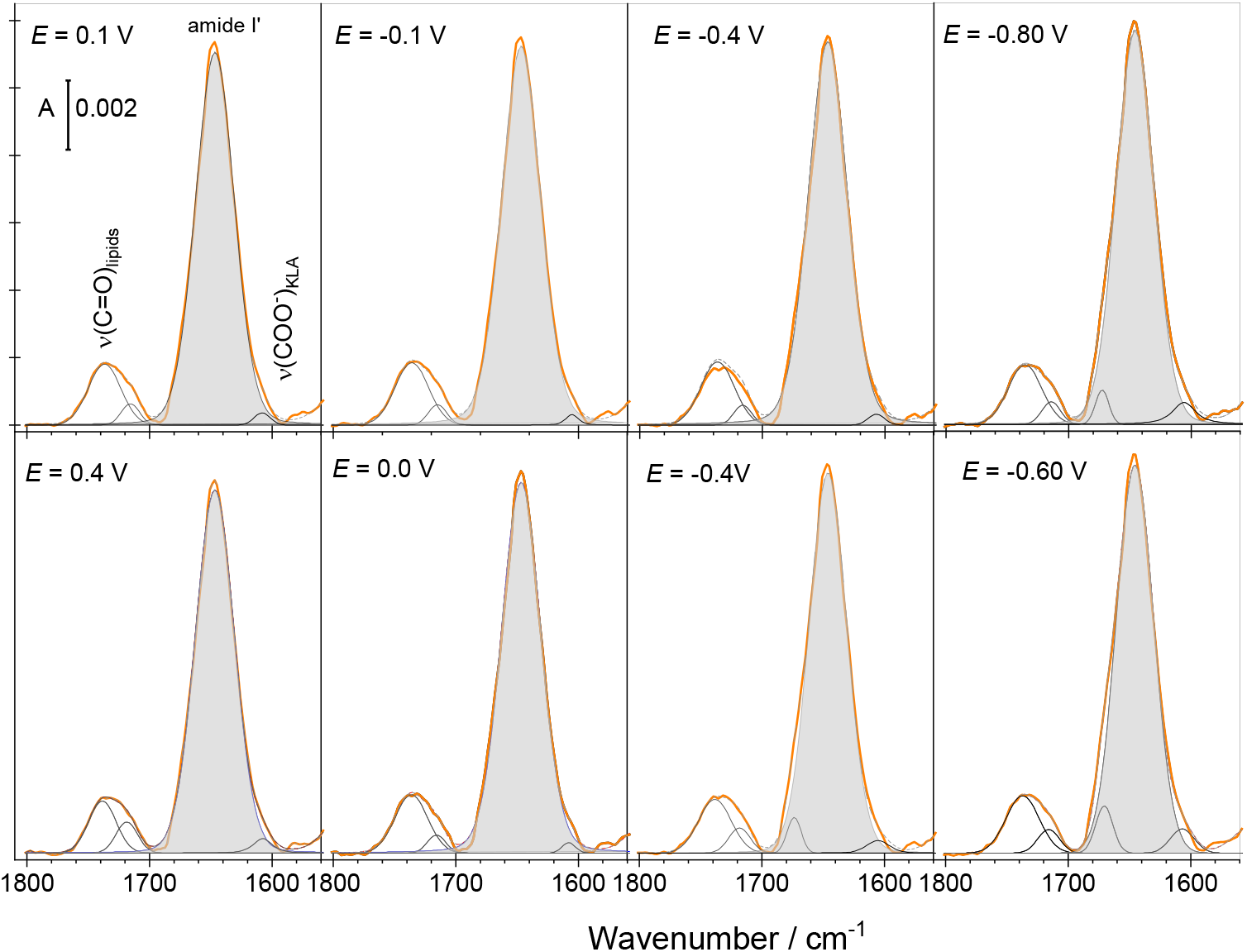
IR spectra of KLA-POPE bilayers on Au(111) after 15 minutes of interaction with melittin (orange lines) recorded at different electrode potentials. The upper and lower panels show results for the negative and positive scans, respectively. Thin black lines show the band deconvolution results. The bands highlighted in gray show the amide I’ mode of *α*-helices in melittin. Measurements were carried out for 10 μM melittin in 50 mM KClO_4_ and

At *E* < *−*0.4 V a weak amide I’ band at 1670-1673 cm^*−*1^ appears in the spectra, indicating the presence of other structural elements in the membrane bound melittin or the presence of poorly hydrated peptide fragments. Similar behavior was observed for 1 μM melittin interacting with the OM for 60 minutes (see, Fig. S7). For both melittin concentrations the potential-dependent spectral changes were reversible, indicating that melittin reached a steady-state well-defined orientation within the OM. Observed enhancement of the intensity of the amide I’ band of α-helices in melittin (centered at 1648-1654 cm^*−*1^) and reversibility of the potential-dependent changes in the shape of the entire amide I’ band confirm an anisotropic orientation of the melittin helix with respect to the OM surface.

The integral intensities of the deconvoluted components of the amide I’ band were used to calculate the average tilt of the long axis of the α-helix in melittin, see section S10. Results of these calculations are shown in Fig. 8. After 15 minutes of 1 μM melittin association with the OM one observes the average tilt angle of 37 *°* ± 5 *°*for the long axis of the peptide helix relatively to the membrane surface normal. MD simulations reveal a similar value of 36° - 39°, as demonstrated above. Both, experimental and simulation results (see Fig. 8A) indicate a tilted arrangement of the helical melittin fragment with respect to the OM surface. However, longer incubation time or higher peptide concentrations induce a reorientation of melittin, see Fig. 8 B,C.

**Figure 8.**
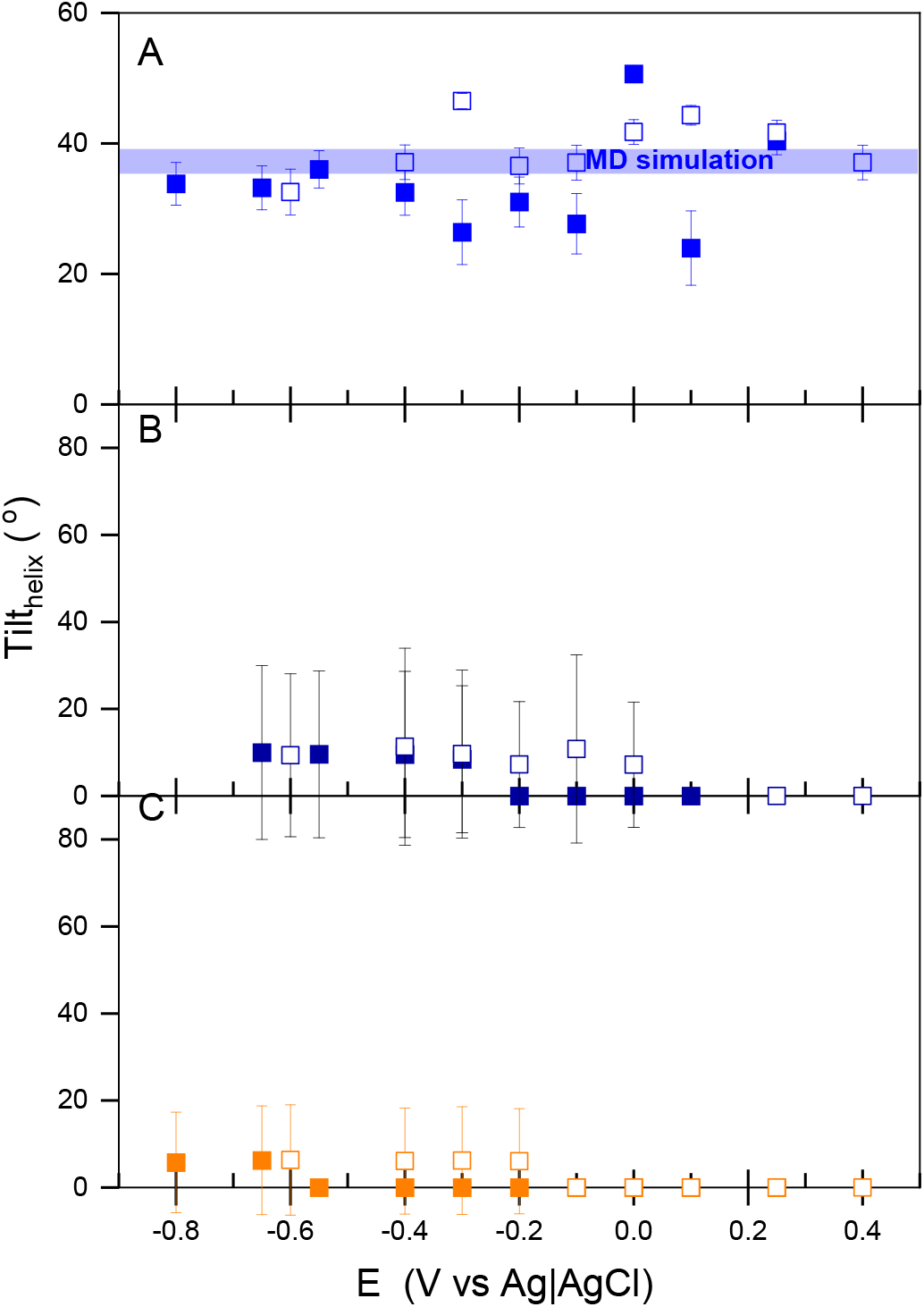
Electrode potential dependence of the tilt angle of the long helix axis in melittin with respect to the KLA-POPE bilayer surface normal. The measurements were done for different melittin concentrations and incubation times as: **A**: 1 μM for 15 min; **B**: 1 μM for 60 min and **C:** 10 μM for 15 min. The electrolyte solution contained 50 mM KClO_4_ and 5 mM Mg(ClO_4_)_2_ in D_2_O. Full and open symbols indicate the tilt angles determined from negative and positive potential scans, respectively. Blue line in panel A shows the tilt of the helix in melittin obtained from the MD simulations.

It is worth mentioning, that at positive potentials, the long axis in the α-helices in melittin is perpendicular to the bilayer surface (the average tilt angle equals 0 *°*). A negative potential shift leads to an increase of the average tilt angle of melittin helices to 10*°* - 15*°*. The potential-dependent change of the helix tilt is related to the emergence of the second amide I’ band at 1670 cm^*−*1^ (see Fig. 7 and Fig. S7); the latter effect is related to the poorly hydrated carbonyl groups and disordered melittin fragments. Earlier investigations showed that in the OM at negative membrane potentials the dehydration of the polar head groups in lipids leads to a reorientation of their hydrocarbon chains.^50^ The membrane associated melittin follows the electric potential driven movement of lipid molecules in the OM.

To evaluate if melittin has an effect on the lipid molecules in both leaflets, *d*_31_-POPE lipid with per-deuterated palmitoyl chain was used to fabricate the inner leaflet. The CD stretching modes appear in the 2215-2070 cm^*−*1^ and are separated by ca. 700 cm^*−*1^ from the CH modes in the KLA and POPE lipids. Due to the conformational disorder of the acyl chains in *d*_31_-POPE and KLA lipids in the liquid disordered OM, the exact tilt angle of the hydrocarbon chains cannot be determined.^81^ The quantitative analysis of some IR absorption modes may become complex due to the overlap of two factors: the conformational disorder of liquid chains and molecular disorder caused by different orientations of an adsorbed molecule. The order parameter of the methylene groups depends on both types of disorder.^82^ The order parameter of the deuterated acyl chains (*S*_CD_) in the pure KLA-*d*_31_-POPE bilayer, as well as the *S*_CD_ for the bilayer exposed to 10 μM melittin for 15 minutes were calculated and are shown in Fig. 9.

**Figure 9.**
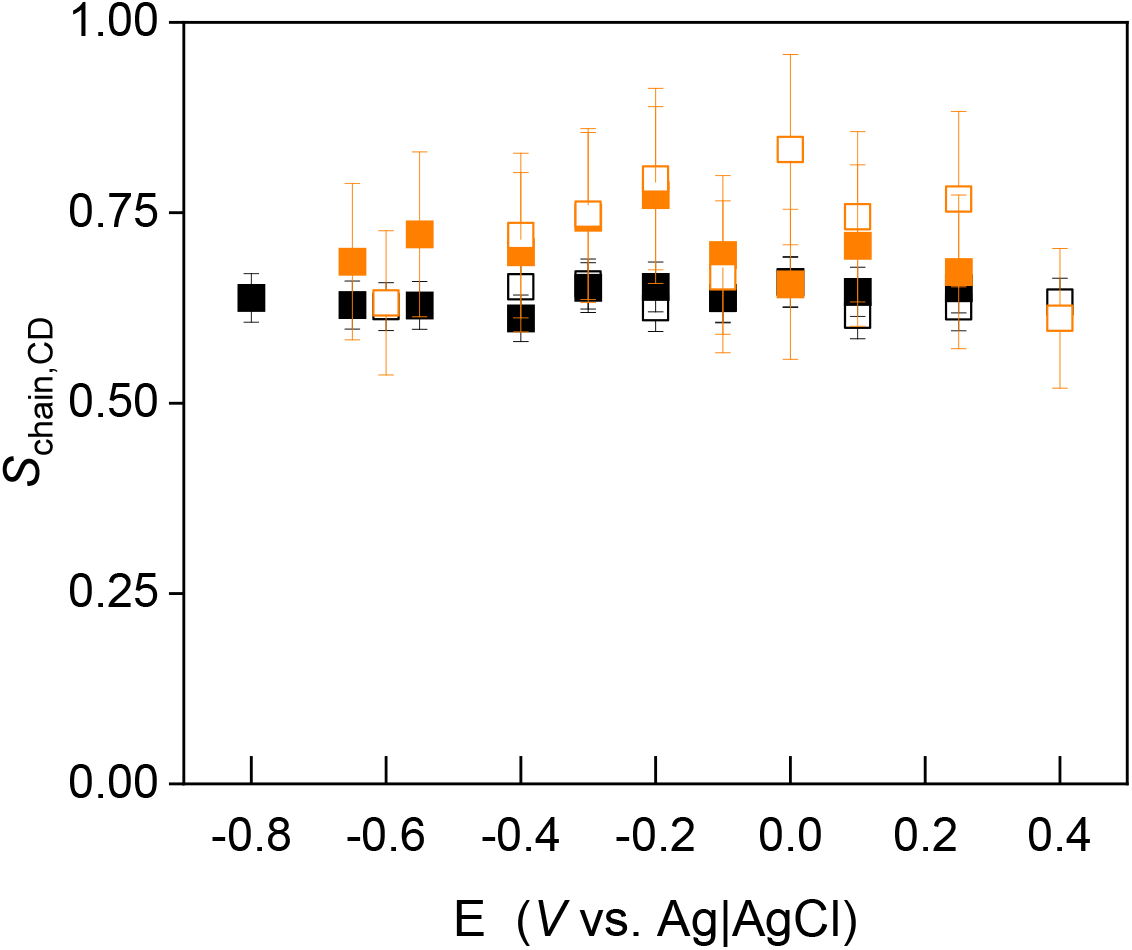
The order parameter (*S*_*CD*_) of the per-deuterated palmitoyl chain in the KLA-*d*_31_-POPE bilayer (black squares) and the order parameter for the bilayer exposed to 10 μM melittin for 15 minutes (orange squares). The electrolyte solution contained 50 mM KClO_4_ and 5 mM Mg(ClO_4_)_2_ in H_2_O. Full and open symbols indicate the order parameter values determined from negative and positive potential scans, respectively.

**Figure 10.**
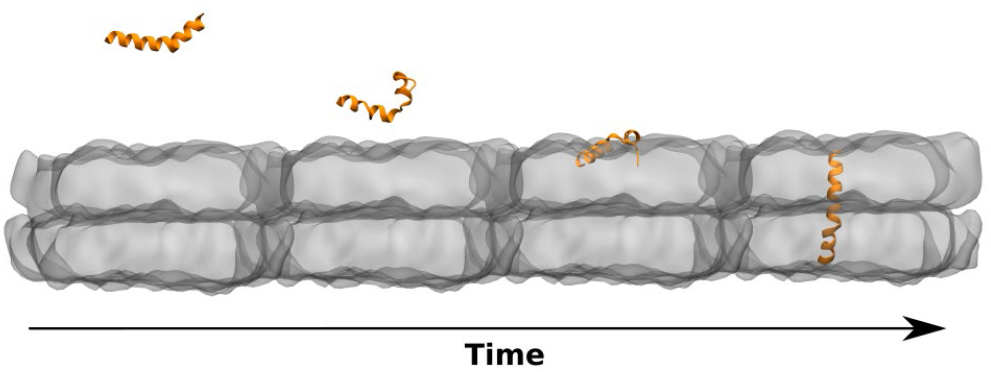
For Table of Contents Only

Binding of melittin to the OM leads to a small increase in *S*_CD_, indicating that the peptide interacts with the hydrophobic fragment of the *d*_31_-POPE in the inner leaflet. The increase in the *S*_CD_, as well as previously reported increase in the *S*_CH_ in the KLA-POPE bilayer^73^, indicate an ordering effect of melittin on the hydrophobic part of the OM. Thus, melittin insertion into the hydrophobic membrane region induces pores and/or channel formation in the bilayer that involve rearrangements of the KLA and POPE, leading to an improved packing of the lipids in the membrane.

## Conclusion

MD simulations and PM IRRAS with electrochemical control reveal molecular scale changes accompanying melittin interacting with a model OM. The interaction begins with a rapid formation of hydrogen bonds between the positively charged Lys21, Arg22, Lys23 and Arg24 residues at the C-terminus in melittin and hydroxyl, carboxylate and phosphate residues in KLA in the outer leaflet of the OM. Formation of these hydrogen bonds immobilizes the α-helical melittin fragment in the polar region of the outer leaflet. In contrast, the N-terminus has a poorly defined structure and displays a large degree of flexibility. MD simulations reveal melittin penetration into the interfacial hydrophilic-hydrophobic region in the lipid A part of KLA. The simulations indicate an increase in the helical content in melittin secondary structure upon the peptide binding to the membrane; this conclusion is in a good agreement with the experimental PM IRRAS results. Here, melittin interacting with the OM adopts predominantly the α-helical conformation with elapsing time. Such a conformational change occurs in melittin upon its insertion into the hydrophobic part of the OM.^3^ Adaptation of a helical conformation in melittin leads to its reorientation within the membrane; the helices align preferentially perpendicular to the membrane surface making it possible for the flexible N terminal to anchor the peptide deeply into the acyl chains region of the OM. Spectroscopic evidence supporting this rationale follows from the isotopic substitution of POPE in the inner leaflet of the OM, where the average packing of the acyl chains is improved not only in KLA^80^, but also in POPE. Thus, melittin has the ability to migrate from the polar region of the outer leaflet deep into the hydrophobic region of the inner leaflet in the OM.

Specific chemical groups in melittin and KLA directly involved in the peptide membrane association could be established through IRS. Interaction between melittin and the negatively charged carboxylate group in KLA was observed as a down-shift of the *ν*_as_(COO^*−*^) stretching mode in KLA, caused by a substitution of Mg^2+^ ions by an amide group of melittin. The removal of divalent ions from the saccharide fragment of lipopolysaccharides destabilizes the OM ^31,32^ and aligns with the observed increase in the membrane capacitance. On the other hand, the reorientation of α-helices in membrane bound melittin also contribute to an increase in the measured capacitance.

Despite increase in the measured membrane capacitance, in the melittin bound bilayer the acyl chains have more compact packing indicating that the peptide penetration into the OM does not dissolve the membrane. The effect could readily be explained by the exceptional conformational flexibility of melittin as could be revealed by a tandem approach relying on spectroscopic measurements and atomistic MD simulations. The investigated melittin serves as a model case study for the AMPs interaction with the cell envelop of Gram-negative bacteria. The mechanism deciphered here turns out to be vastly different, and maybe even unique, in comparison to the previously known AMPs action on phospholipid bilayers.

## Supporting Information document is available

Supporting information contains the following sections: Formation of the asymmetric models outer membrane of Gram-negative bacteria, Area per lipid of the outer membrane and volume of the simulation box during the equilibration simulation, Possible orientation of melittin on the membrane surface and their effect on the capacitance of a model membrane deposited on an electrode surface, Calculation of electrostatic and van der Waals contribution to the interaction energy, Secondary structure analysis of melittin interacting with the outer membrane, Location and conformation of melittin associated with the KLA-POPE bilayer, Deconvolution of the amide I’ mode in melittin, PM IRRA spectra of the KLA-POPE bilayer after interaction with 1 μM melittin, and Determination of the helix tilt angle from the PM IRRA spectra.

## Author Contributions

BK performed all the experiments, analyzed the electrochemical data, made background corrections of the PM IRRA spectra. IB performed the quantitative analysis of the PM IRRA spectra. JCS performed all calculations. JCS and LG performed quantitative analysis of the calculation results. The manuscript was written by IB, IAS, JCS and LG. The final version of the manuscript was approved by the authors. IB, IAS supervised the project. IB, IAS acquired funding and resources for this project.

## Acknowledgements

The authors would like to thank the Volkswagen Foundation (Lichtenberg professorship awarded to I.A.S.), the Deutsche Forschungsgemeinschaft (SFB 1372 Magnetoreception and Navigation in Vertebrates, no. 395940726, and HYP*MOL – Hyperpolarization in molecular systems (TRR386/1 - 2023, no. 514664767 to I.A.S. BR-3961/4 to IB), the Ministry for Science and Culture of Lower Saxony “Simulations Meet Experiments on the Nanoscale: Opening up the Quantum World to Artificial Intelligence (SMART)” and “Dynamik auf der Nanoskala: Von kohärenten Elementarprozessen zur Funktionalität (DyNano)”. Computational resources for the simulations were provided by the CARL Cluster at the Carl-von-Ossietzky University, Oldenburg, supported by the DFG and the Ministry for Science and Culture of Lower Saxony. The authors also gratefully acknowledge the computing time granted by the Resource Allocation Board and provided on the super-computer Lise and Emmy at NHR@ZIB and NHR@ Göttingen as part of the NHR infrastructure. The calculations for this research were conducted with computing resources under the project nip00058.

