## Supporting Information (SI) for "How do antimicrobial peptides interact with the outer membrane of Gram-negative bacteria? Role of lipopolysaccharides in the peptide binding, anchoring and penetration"

Justus C. Stephani[1], Luca Gerhards[1], Bishoy Khairalla[2,5], Ilia A. Solov'yov[1,3,4]*, and Izabella Brand[2]*

[1] *Institute of Physics, Carl von Ossietzky University of Oldenburg, 26111 Oldenburg, Germany*

[2] *Department of Chemistry, Carl von Ossietzky University of Oldenburg, 26111 Oldenburg, Germany*

[3] *Research Center Neurosensory Science, Carl von Ossietzky University of Oldenburg, 26111 Oldenburg, Germany*

[4] *CeNaD – Center for Nanoscale Dynamics, Carl von Ossietzky University of Oldenburg, 26111 Oldenburg, Germany*

[5] *Current address: Friedrich-Alexander-Universität Erlangen-Nürnberg, Department of Biology, Pharmaceutical Biology, Staudtstr. 5, 91058 Erlangen, Germany*

### S1. Formation of the asymmetric model outer membrane of Gram-negative bacteria

Langmuir-Blodgett (LB) and Langmuir-Schaefer (LS) transfers were used to prepare asymmetric KLA-POPE bilayers on a gold surface. First, a POPE (or $d_{31}$-POPE) monolayer was transferred from the aqueous subphase by a vertical LB withdrawal, see Fig. S1A. Withdrawal of a hydrophilic gold substrate from the aqueous subphase through the air|water interface covered by the phospholipid monolayer gave the inner leaflet of the model outer membrane. Next, a monolayer of KLA on a 0.1 M $KClO_4$ and 5 mM $Mg(ClO_4)_2$ aqueous subphase was compressed to the surface pressure $\Pi = 30$ mN m$^{-1}$ and a horizontal LS transfer was used to fabricate the second leaflet of the model outer membrane, see Fig. S1B. During this transfer, the hydrophobic surface of the POPE modified gold surface and was exposed to the hydrophobic hydrocarbon chains in the KLA monolayer present at the aqueous electrolyte|air interface. The gold substrate was covered by a Y-type lipid bilayer as shown schematically in Fig. S1C.

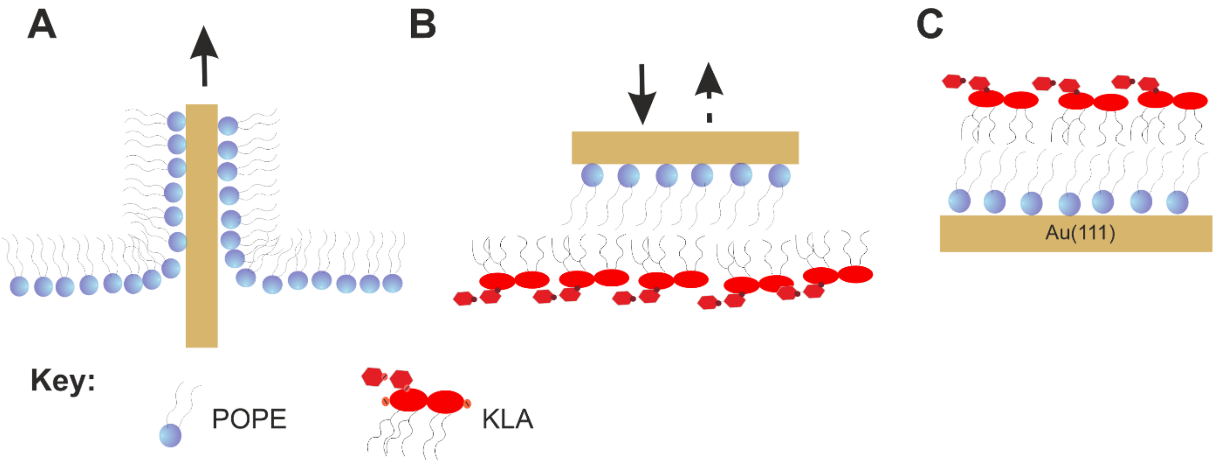


**Figure S1.** Illustration of the fabrication procedure of the asymmetric KLA-POPE model outer membrane of Gram-negative bacteria **A**: transfer of the inner POPE leaflet by Langmuir-Blodgett vertical withdrawal, **B**: transfer of the outer KLA leaflet by Langmuir-Schaefer method and **C**: molecular scale order in the LB-LS transferred bilayer.

**S2. Area per lipid of the outer membrane and volume of the simulation box during the equilibration simulation**

During the equilibration process of the model outer membrane, the volume of the simulation box and the average area per lipid attributed to KLA and POPE molecules decreased. Figure S2A shows the average area per KLA lipid in the outer leaflet of the membrane over simulation time, while Fig. S2B shows the average area per POPE phospholipid in the inner leaflet of the outer membrane. The average areas per lipid $A_{KLA}$ and $A_{POPE}$ converge to the values 1.87 nm$^2$ and 0.58 nm$^2$, respectively. These values agree well with the LB-LS transfer conditions at with the average area per $A_{KLA}$ was 1.96 nm$^2$ and $A_{POPE}$ – 0.67 nm$^2$.[1] The time evolution of the total volume of the simulation box $V_S$ is shown in Fig. S2C.

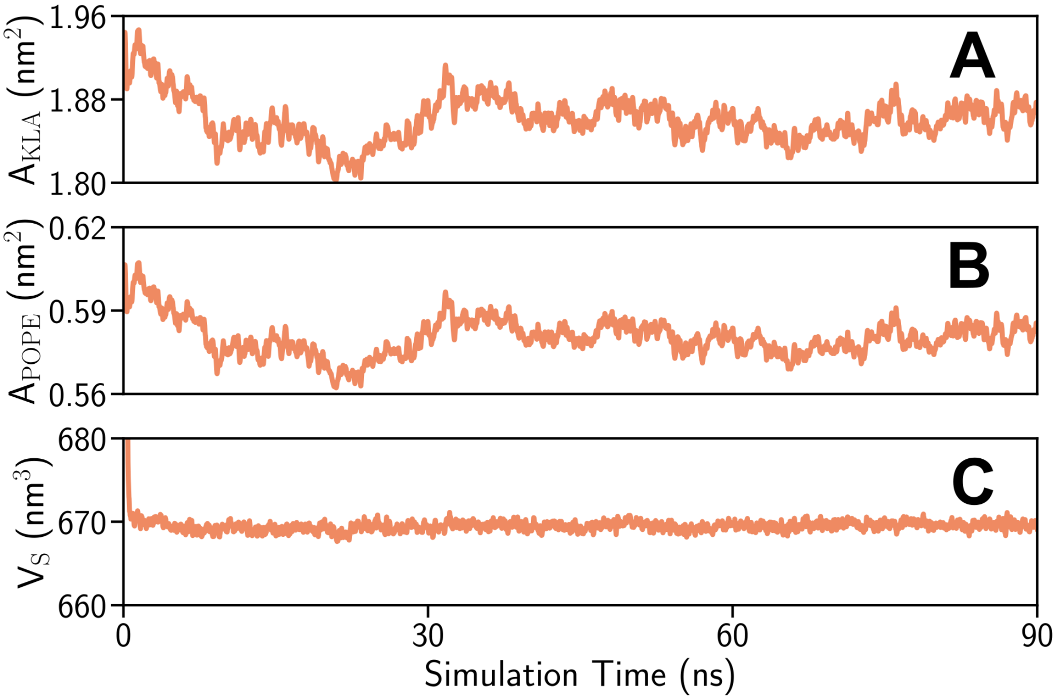


**Figure S2. A**: Time evolution of the average area per KLA lipid in the outer leaflet of the model outer membrane; **B**: Time evolution of the average area per POPE phospholipid on the inner leaflet of the model outer membrane. The values were computed as an average over all lipids present in the simulation box. **C**: Volume of the simulation box $V_S$ over simulation time.

**S3. Possible arrangements of melittin on the membrane surface and their effect on the capacitance of a model membrane deposited on an electrode surface**

Figure S3 shows possible arrangements of melittin in the model outer membrane of Gram-negative bacteria.

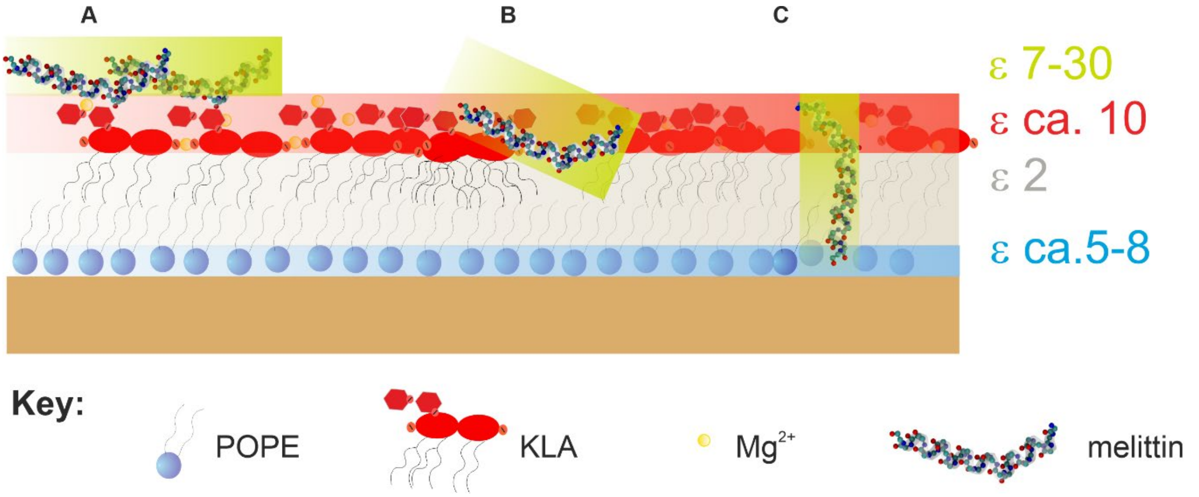


**Figure S3**. Schematic structure of the KLA-POPE model outer membrane on a gold electrode surface with possible arrangements of melittin interacting with the membrane from solution **A**: parallel **B**: tilted and **C**: perpendicular orientations. The dielectric constants of polar head groups and hydrocarbon chains in lipids as well as of the peptide are given in the figure.

The capacitance of a film modifying an electrode surface is

$$C = \frac{\varepsilon_0 \varepsilon A}{d} \quad , \qquad \text{(S1)}$$

where $\varepsilon_0$ is the permeability of vacuum, $\varepsilon$ is the dielectric constant of the film at the electrode surface, $d$ is the thickness of the film, and $A$ is the surface area of the electrode. The only purpose of presenting this equation is due to the description of the dependence of the measured capacitance on the dielectric constants of molecules present in the film.

**S4. Calculation of electrostatic and van der Waals contribution to the interaction energy**

The electrostatic contribution to the interaction energy of a residue $x$ in the peptide with the membrane is calculated as:

$$U_{\text{elec}}(x) = \frac{1}{4\pi\varepsilon_0}\Sigma_{(i=1)}^{N}\Sigma_{(j=1)}^{M}\frac{q_i q_j}{r_{ij}} \ , \tag{S2}$$

where $r_{ij}$ is the distance between two charges $q_i$ and $q_j$, where the first summation goes over all the $N$ atoms of the residue of interest, while the second summation goes over all the $M$ atoms of the membrane. The van der Waals contribution to the interaction energy could be calculated using the Lennard-Jones potential as

$$U_{vdW}(x) = \sum_{i=1}^{N}\sum_{j=1}^{M}\varepsilon_{ij}\left[\left(\frac{\sigma_{ij}}{r_{ij}}\right)^{12} - 2\left(\frac{\sigma_{ij}}{r_{ij}}\right)^{6}\right], \tag{S3}$$

where $r_{ij}$ is the distance between two atoms $i$ and $j$, where the first summation goes over all the $N$ atoms of the residue of interest, while the second summation goes over all the $M$ atoms of the membrane. $\varepsilon_{ij}$ and $\sigma_{ij}$ are the equilibrium van der Waals energy and distance for a given pair of atoms.

**S5. Secondary structure analysis of melittin interacting with the outer membrane**

The secondary structure of the peptide was determined using the STRIDE algorithm in VMD.[2] Helicity is defined as the amount of α-, $3_{10}$- and π-helices in the secondary structure of the peptide. The time dependency of the helicity is plotted in Fig. S4A for the three production simulations of the composite systems (Sim. 1 - Sim. 3) and the control simulation of melittin in water (Control). The helicity value decreased most notably in the Control simulation from 75% to 35% during the simulation time of 200 ns, while the helicity stayed largely constant in Sim. 1 and slightly decreased in Sim. 2 and Sim. 3. The results indicate that the interaction of the peptide with the membrane stabilizes the helical structures.

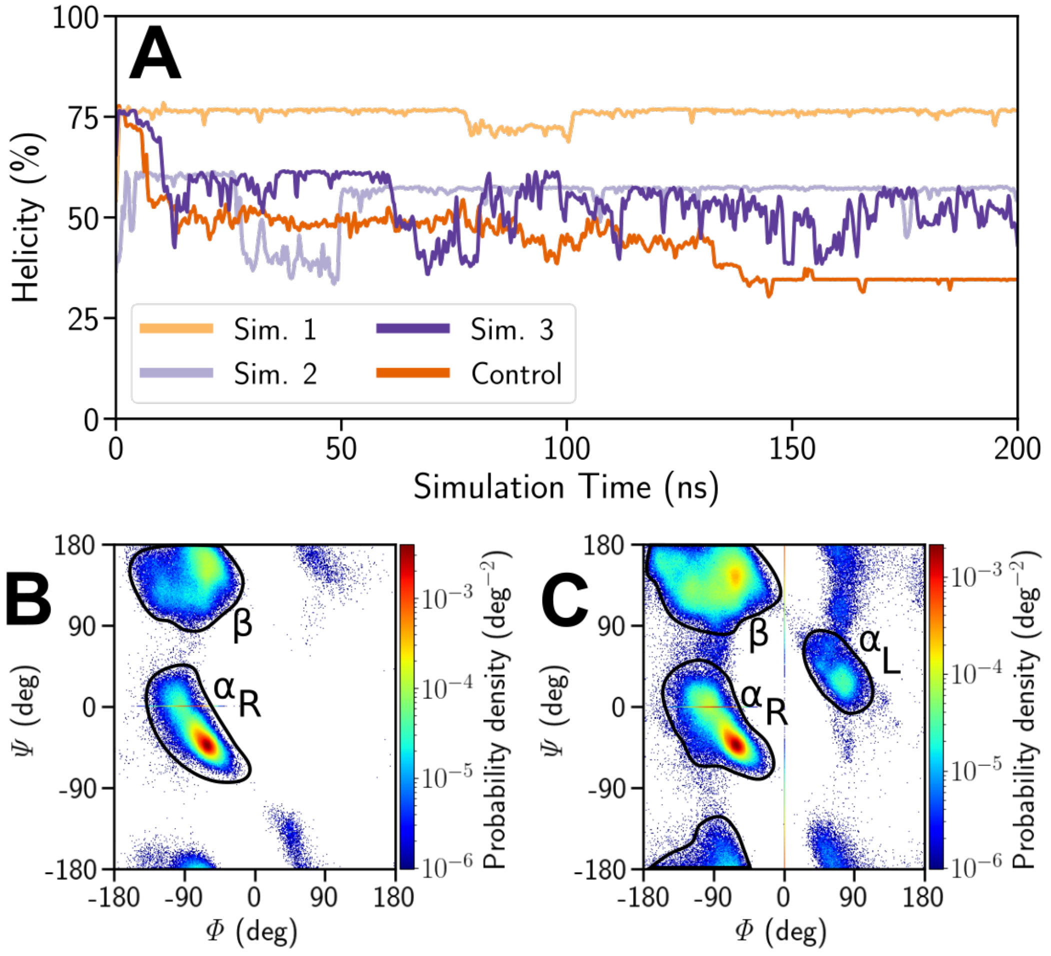


**Figure S4. A**: Time dependency of helicity of melittin (percent) for the four simulations Sim. 1, Sim. 2, Sim. 3 and Control. **B/C**: Probability distributions of the backbone dihedral angles $\psi$ and $\varphi$ computed for Sim. 1 (**B**) and Control (**C**). The regions that correspond to dominant peptide conformations are marked. The white area corresponds to the sterically forbidden conformations of the peptide.

Figure S4B-C show the Ramachandran plots of the probability distribution of the backbone dihedral angles $\psi$ and $\varphi$ computed for Sim. 1 and Control, respectively. While in Sim. 1 the right turning α-helix is the dominant secondary structure followed by the β-sheet, in the Control simulation the probability to observe α-helix decreases and other conformations are possible, further indicating the unfolding of the α-helix of melittin in solution.

### S6. Location and conformation of melittin associated with the KLA-POPE bilayer

Figure S5 shows the configuration of melittin associated with the membrane after 200 ns of the three simulations.

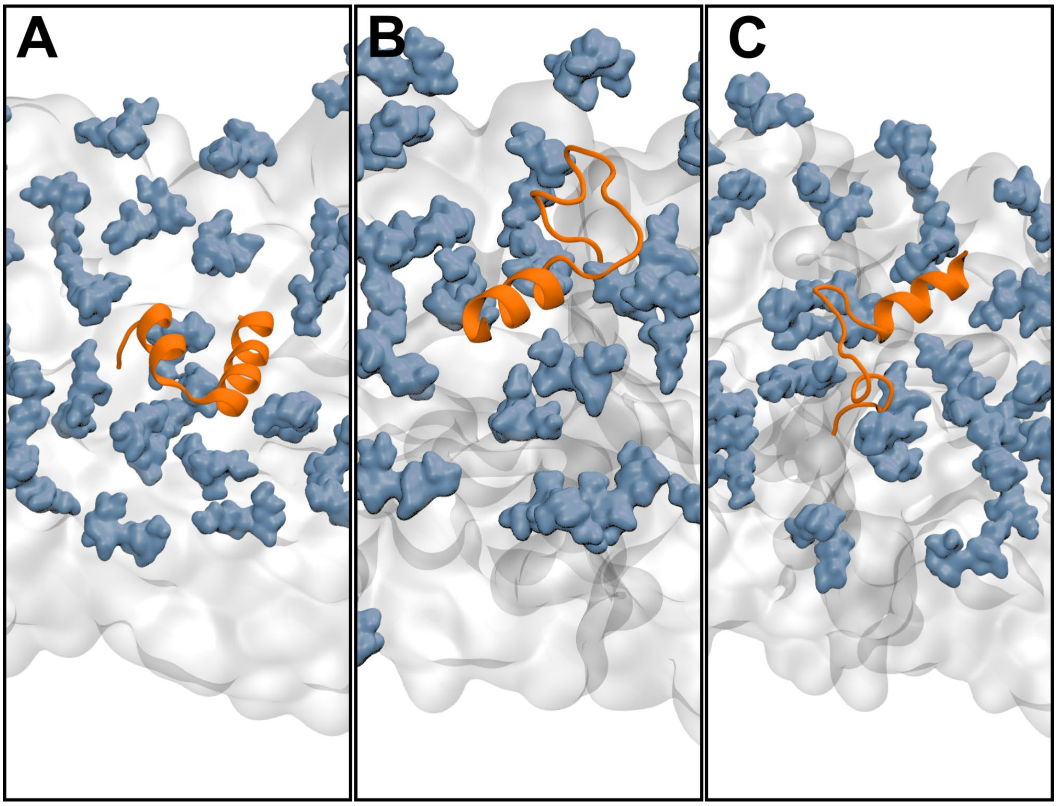


**Figure S5.** Characteristic configurations of melittin (orange) bound to the model outer membrane in **A**: Sim. 1; **B**: Sim. 2 and **C**: Sim. 3. The upper leaflet of the outer membrane is shown in silver while the saccharide residues in KLA are highlighted in blue.

### S7. Deconvolution of the amide I’ band in melittin

The second derivative and Fourier self-deconvolution were used to deconvolute the measured IR spectra. Fourier self-deconvolution was done to fit the measured amide I’ band with Gauss shaped curves with the full width at half maximum set to 23 $cm^{-1}$. The wavenumbers of the second derivative minima overlapped with the position of the maxima of the Fourier self-deconvoluted bands yielding the number of the deconvoluted bands. The measured IR spectra were deconvoluted using OPUS software (Bruker, Germany) by setting the number of amide I’ band components and their positions (wavenumbers) to the results of the second derivative and Fourier self-deconvolution. The individual components of the amide I’ band were fitted with Gauss curves. Figure S6 shows the deconvoluted measured IR spectra (bottom panels), the Fourier self-deconvoluted spectra (bottom panels, gray lines), and the second derivatives of the experimental spectra (top panel) of the KLA-POPE lipids upon interaction with melittin.

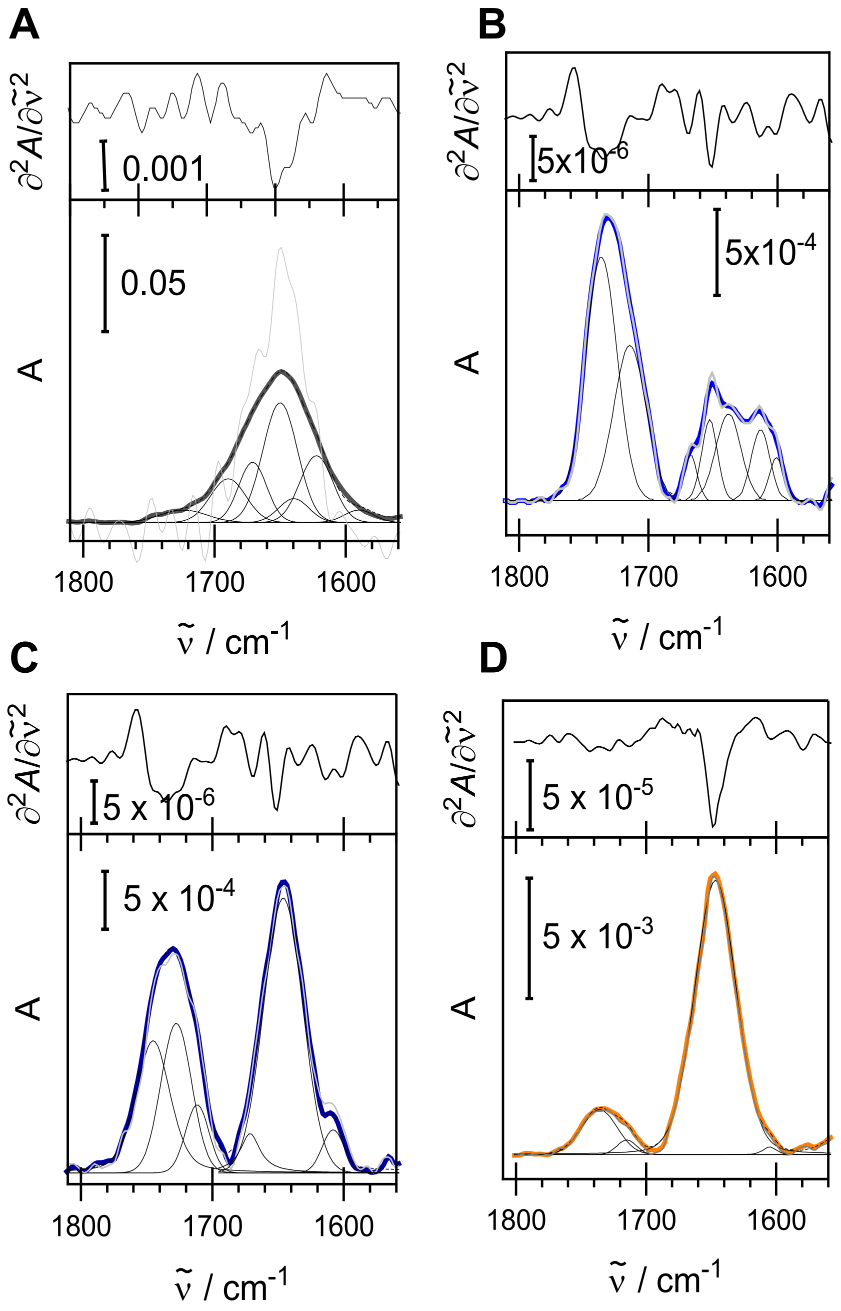


**Figure S6**. Upper panel: the second derivative of the measured IR spectra and the bottom panel: corresponding IR spectra (thick color lines) KLA-POPE lipid systems after interaction with melittin. Thin black lines show the band deconvolution results. **A**: attenuated total reflection IR spectrum of 0.44 mM melittin after 1h of interaction with KLA-POPE vesicles, **C-D**: PM IRRA IR spectra of the KLA-POPE bilayer at $E = 0.0$ V vs Ag|AgCl after **B**: 15 min interaction with1 μM melittin, **C**: 1h interaction with 1μM melittin and **D**: 15 min interaction with 10 μM melittin. All spectra were recorded in 50 mM $KClO_4$ and 5 mM $Mg(ClO_4)_2$ in $D_2O$. The scale bars show absorbance in arbitrary units.

### S8. PM IRRA spectra of the KLA-POPE bilayer after interaction with 1 μM melittin

Figure S7 shows the PM IRRA spectra of the KLA-POPE bilayer exposed to 1 μM melittin solution for 60 minutes.

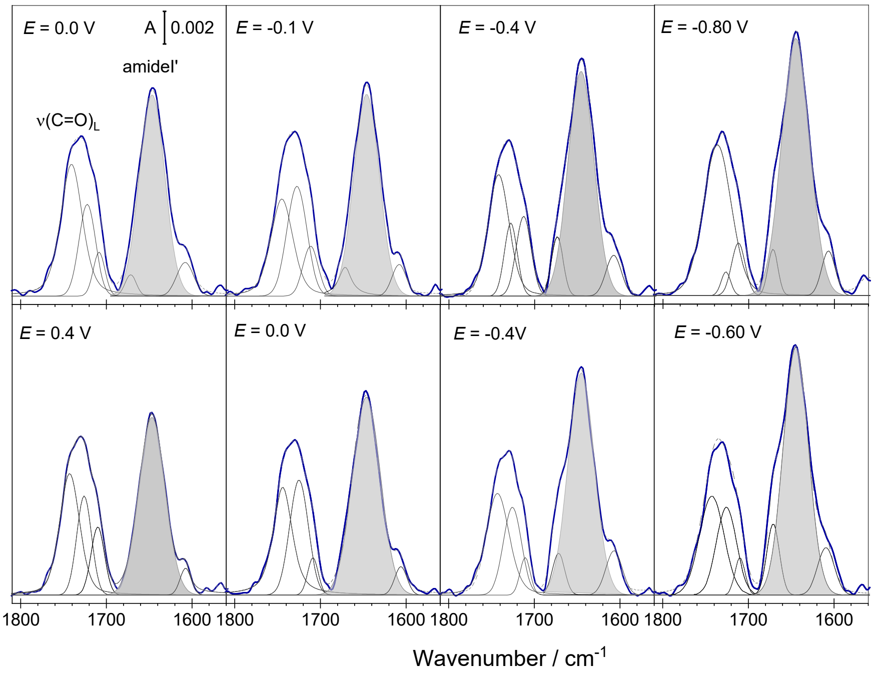


**Figure S7.** IR spectra of KLA-POPE bilayers on Au(111) after 1 h of interaction with 1 μM melittin (dark blue lines) recorded at different electrode potentials. The upper and lower panels show results for the negative and positive scans, respectively. Thin black lines show the band deconvolution results. The bands highlighted in gray show the amide I' mode of α-helices in melittin. Measurements were carried out in 50 mM $KClO_4$ and 5 mM $Mg(ClO_4)_2$ in $D_2O$. The absorbance is shown in arbitrary units.

### S9. Determination of the helix tilt angle from the PM IRRA spectrum

According to the surface selection rule of IRRAS[3, 4], in an anisotropic film, the intensity of an IR absorption mode depends on the surface concentration of species adsorbed on a solid surface and on the average orientation of a given transition dipole vector $\left(\vec{\mu}\right)$ vs. electric field $\left(\vec{E}\right)$ (surface normal), characterized by the angle $\theta$. In a PM IRRA experiment an average orientation of a given functional group can be calculated form the $\langle\theta\rangle$ value. Depending on the orientation of the molecule in a film, some IR absorption bands are enhanced while others attenuated in the IRRA spectrum. Figure S8 shows two limiting cases for the orientation of an α-helical fragment in a peptide adsorbed on a solid surface leading to the enhancement and cancellation of the amide I' band intensity in the IRRA spectrum. A parallel orientation of the $\vec{\mu}$ and $\vec{E}$ vectors causes their strong coupling enhancing the intensity of the IR absorption band of the amide I' band (Fig. S8A).

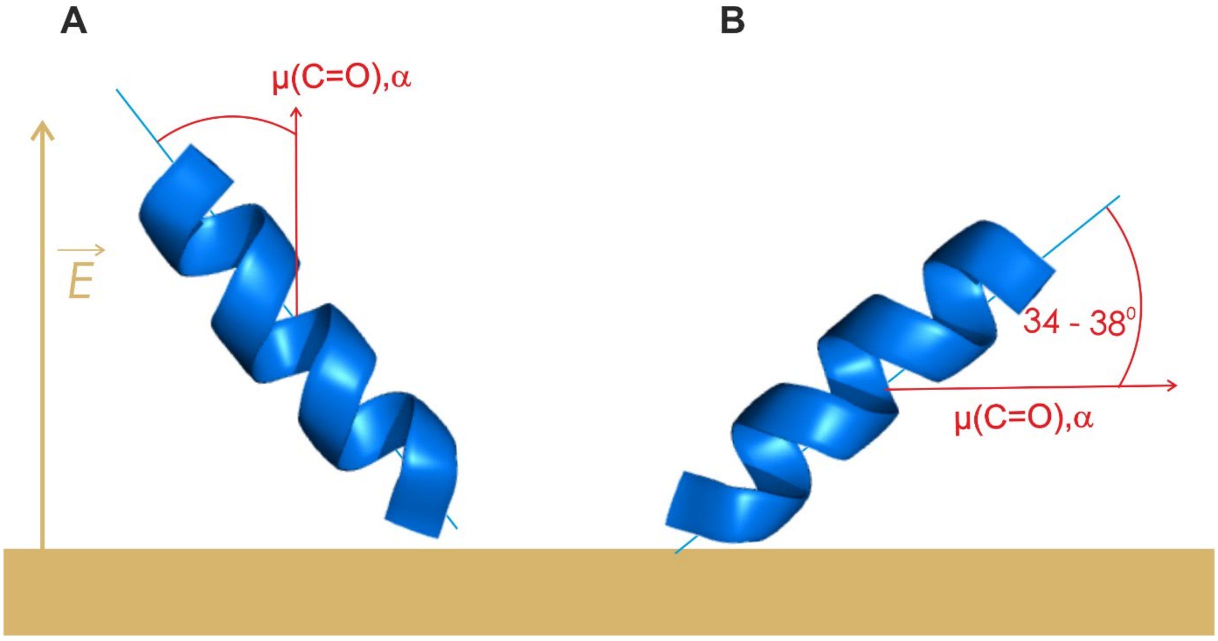


**Figure S8.** Limiting cases for the orientation of an α-helical peptide fragment adsorbed on a solid surface in which the transition dipole moment of the amide I' band is **A**: parallel and **B**: normal to the direction of the electric field vector of the reflected IR radiation. The blue lines show the direction of the long axis of the α-helix. The direction of the transition dipole vector of the amide I' mode (red arrow) and the direction of the electric field vector of the *p*-polarized light at the phase boundary (gold arrow) are shown in the figure.

Once the angle between the $\vec{\mu}$ and $\vec{E}$ vectors equals 90 º (Fig. S8B) the integral intensity of the amide I' band equals zero. In this case there is no coupling of the transition dipole and the electric field vectors.

For a peptide where the α-helical fraction is known, one can calculate the $\langle\theta\rangle$ between the transition dipole moment $\vec{\mu}$ of the deconvoluted amide I' band of α-helices relatively to the surface normal, being colinear with the direction of the electric field vector

$$\cos^2\langle\theta\rangle = \frac{1}{3}\frac{c_{Ex}}{c_{\mathrm{R}}}. \tag{S4}$$

Here $c_{\mathrm{Ex}}$ corresponds to the percent content of the amide I'α-helix band in the entire amide I' band in a measured PM IRRA spectrum and the $c_{\mathrm{R}}$ is the percent content of the α-helical structures in the solution phase (random distribution). The angle $\langle\theta\rangle$ can be used to calculate the order parameter $S$ as follows[5]

$$S = \left\{\frac{1}{2}\left(3\left(\cos^2\langle\theta\rangle\right)-1\right)\right\}. \tag{S5}$$

Finally, the order parameter of the long axis of the α-helix ($S_{\mathrm{helix}}$) can be calculated as

$$S_{helix} = \frac{2S}{3\cos^2\alpha - 1}, \tag{S6}$$

where α is the angle between the long axis of the α-helix and the transition dipole moment of the amide I' band of α-helices. In an α-helical protein fragment the transition dipole vector of the amide I' mode $\vec{\mu}$ makes an angle of 34 ° – 38 ° vs. the long axis of the α-helix[6, 7], see Fig. S8. The order parameter $S_{\mathrm{helix}}$ was used to calculate the tilt of the helix with respect to the surface normal ($\mathrm{Tilt_{helix}}$).
